# PI3Kδ hyper-activation promotes the development of B cells that exacerbate *Streptococcus pneumoniae* infection in an antibody-independent manner

**DOI:** 10.1101/280651

**Authors:** Anne-Katrien Stark, Anita Chandra, Krishnendu Chakraborty, Rafeah Alam, Valentina Carbonaro, Jonathan Clark, Srividya Sriskantharajah, Glyn Bradley, Alex G. Richter, Edward Banham-Hall, Menna R. Clatworthy, Sergey Nejentsev, J. Nicole Hamblin, Edith M. Hessel, Alison M. Condliffe, Klaus Okkenhaug

**Author notes:** Joint first authors (Anne-Katrien Stark, Anita Chandra).

## Abstract

*Streptococcus pneumoniae* is a major cause of pneumonia and a leading cause of death world-wide. Antibody-mediated immune responses can offer protection against repeated exposure to *S. pneumoniae*, yet vaccines only offer partial protection. Patients with Activated PI3Kδ Syndrome (APDS) are highly susceptible to *S. pneumoniae*. We generated a conditional knockin mouse model of this disease and identified a CD19^+^B220^−^ B cell subset that is induced by PI3Kδ signaling, is resident in the lungs, and which promotes increased susceptibility to *S. pneumoniae* during the early phase of infection via an antibody-independent mechanism. We show that an inhaled PI3Kδ inhibitor improves survival rates following *S. pneumoniae* infection in wild-type mice and in mice with activated PI3Kδ. These results suggest that a subset of B cells in the lung can promote the severity of *S. pneumoniae* infection, representing a novel therapeutic target.

## Introduction

*Streptococcus pneumoniae* is an invasive extracellular bacterial pathogen and is a leading cause of morbidity and mortality. Although *S. pneumoniae* can cause disease in immunocompetent adults, it commonly colonizes the upper airways without causing disease. The World Health Organization has estimated that there are 14.5 million episodes of severe pneumococcal disease and that 1.6 million people die of pneumococcal disease every year^1^. Despite the implementation of global vaccination programs, *S. pneumoniae* infection remains a major disease burden^2–4^.

Invasive *S. pneumoniae* infection is a major cause of lower airway infections (pneumonia), sepsis and meningitis. Healthy people at the extremes of age are more susceptible to pneumococcal disease, as are people with chronic obstructive pulmonary disease (COPD), however those at greatest risk are patients with splenic dysfunction or immune deficiency. This increased susceptibility results at least in part from the lack of protective antibodies against conserved protein antigens or against polysaccharides that form part of the pneumococcal capsule^5^. Indeed, the protective role of antibodies in pneumococcal disease is most obvious in individuals with congenital (primary) immunodeficiencies (PIDs). This was first recognized in a patient with X-linked agammaglobulinemia (XLA), a syndrome subsequently shown to be caused by a block in B cell development due to loss-of-function mutations in *BTK*^6–8^. These patients remain highly susceptible to *S. pneumoniae* into adulthood, but can be effectively treated by the administration of immunoglobulins from healthy donors.

We and others have recently described cohorts of immune deficient patients with activating mutations in *PIK3CD*, the gene encoding the p110δ catalytic subunit of phosphoinositide 3-kinase δ (PI3Kδ)^9–11^. PI3Kδ is a lipid kinase that catalyzes the phosphorylation of the phosphatidylinositol-(4,5)-bisphosphate lipid to produce phosphatidylinositol-(3,4,5)-trisphosphate (PIP_3_). PI3Kδ is expressed in cells of the immune system and regulates many aspects of immune cell signaling, particularly in lymphocytes^12, 13^. Activated phosphoinositide 3-kinase δ syndrome (APDS) is a combined immunodeficiency affecting T and B cells. APDS patients suffer from recurrent sinopulmonary infections, with *S. pneumoniae* being the most commonly isolated pathogen^14^. 85% of APDS patients have been diagnosed with pneumonia^15^. APDS patients are also more likely to develop structural lung damage (bronchiectasis) than patients with other PIDs^14^. The mechanism underpinning the increased susceptibility to pneumococcal infection in APDS is unclear^12^. Although APDS patients often lack IgG2, the protection afforded by immunoglobulin replacement therapy is not as robust as that observed in patients with pure antibody deficiencies, suggesting that antibody-independent PI3Kδ-driven mechanisms may be involved^14^. The monogenic nature of APDS allows us to dissect mechanisms of susceptibility to *S. pneumoniae* infection on cellular and molecular levels, and to determine whether PI3Kδ inhibitors may help reduce the susceptibility to *S. pneumoniae.* If so, PI3Kδ inhibitors, that are in development for the treatment of inflammatory and autoimmune diseases, might also have wider applications to reduce the pathological consequences of *S. pneumoniae* infection^16^. In this study, we have explored mechanisms by which PI3Kδ hyperactivation drives susceptibility to *S. pneumoniae* infection. We found that the administration of the PI3Kδ-selective inhibitor nemiralisib (GSK-22696557)^17, 18^ reduced the severity of pneumococcal disease in wild-type mice. To investigate this further, we generated a p110δ^E1020K^ mouse model that accurately recapitulates the genetics and immunological phenotype of APDS, and displays increased susceptibility to *S. pneumoniae* infection. We show that this susceptibility segregates with enhanced PI3Kδ signaling in B cells, which exacerbate *S. pneumoniae* infection at early time points before the adaptive immune response comes into play. Of note, we have identified a previously unappreciated population of CD19^+^B220^−^ IL-10-secreting cells that was present in wild-type mice but expanded 10-20 fold in p110δ^E1020K^ mice. We demonstrate that nemiralisib reduces the frequency of IL-10-producing B cells in the lung and improves survival of p110δ^E1020K^ mice. Similarly, a higher proportion of transitional B cells from APDS patients produced IL-10 and this was reduced by nemiralisib. This study provides new insights into the pathogenesis of the early stages of invasive *S. pneumoniae* disease and offers the potential of future therapeutic strategy to alleviate the severity of this disease in susceptible patients.

## Results

### Inhaled nemiralisib alleviates the severity of *S. pneumoniae* infection in mice

Given that APDS patients are more susceptible to *S. pneumoniae*, we sought to determine whether nemiralisib, an inhaled PI3Kδ inhibitor which is in development for the treatment and prevention of COPD excacerbations^17, 18^, would alter susceptibility to airway infections. We treated mice with nemiralisib and then infected them intranasally with *S. pneumoniae.* Nemiralisib-treated mice showed prolonged survival compared to mice given vehicle control (Fig 1). This protection was only effective if the drug was administered before and during infection (Fig 1). By contrast, nemiralisib administration 8h or 24h post infection had no impact on survival of the mice. These data suggest that PI3Kδ modulates the immune response during early *S. pneumoniae* infection, either by inhibiting protective immunity, or by promoting an adverse response.

**Figure 1:**
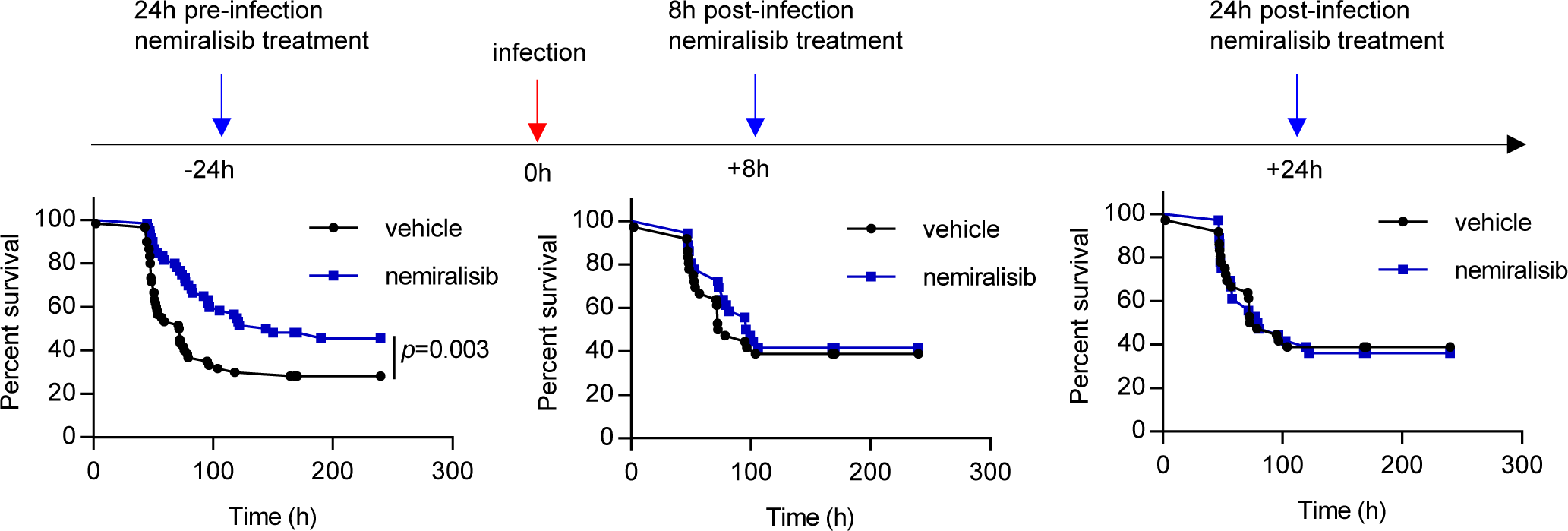
Prophylactic, but not therapeutic treatment with the inhaled PI3Kδ inhibitor nemiralisib mitigates disease severity following *S. pneumoniae* infection in wild-type mice. Wild-type mice were treated twice daily with the inhaled PI3Kδ inhibitor nemiralisib for the duration of the study: when treatment was started 24h prior to infection with *S. pneumoniae*, survival rates were improved. When started 8h or 24h post-infection, the treatment had no effect on survival outcome. (-24h: data from 5 independent experiments combined n=60; +8h/+24h: data from 3 independent experiments combined n=36).

### Mouse model of APDS: hyperactive PI3Kδ signaling and altered B and T cell development in p110δ^E1020K^ mice

We generated a conditional knock-in mouse harboring mutation E1020K in the *Pik3cd* gene that is equivalent to the most common APDS-causing mutation E1021K in humans (Supplementary Fig 1). These mice were subsequently crossed with different Cre-expressing lines to either generate germline mice where p110δ^E1020K^ is expressed in all cells (p110δ^E1020K-GL^) or selectively in B cells using *Mb1*^Cre^ (p110δ^E1020K-B^), in T cells using *Cd4*^Cre^ (p110δ^E1020K-T^) or myeloid cells using *Lyz2*^Cre^ (p110δ^E1020K-M^). We studied p110δ^E1020K^ mice in comparison with wild-type and p110δ^D910A^ mice that have catalytically inactive p110δ^19^.

Initially, we tested if p110δ^E1020K^ mice have increased PI3Kδ activity and display the characteristic immunological phenotype of APDS. Biochemical analyses of B cells and T cells from p110δ^E1020K-GL^ mice confirmed that the kinase is hyperactive (Fig 2). Measurements of PIP_3_ in T cells showed that p110δ^E1020K^ is about 6 times as active as the wild-type kinase following stimulation with anti-CD3 and anti-CD28, but with no evidence for increased basal activity (Fig 2A). In B cells, p110δ^E1020K^ led to increased basal PIP_3_ levels, but it was further increased only about 2-fold compared to wild-type mice after stimulation with anti-IgM (Fig 2B). This pattern resembles results found in patients with APDS^10^. In wild-type and p110δ^E1020K-GL^ cells, the PI3Kδ-selective inhibitor nemiralisib reduced PIP_3_ to the background level observed in p110δ^D910A^ cells, which, as expected, were insensitive to nemiralisib (Fig 2A, B).

**Figure 2:**
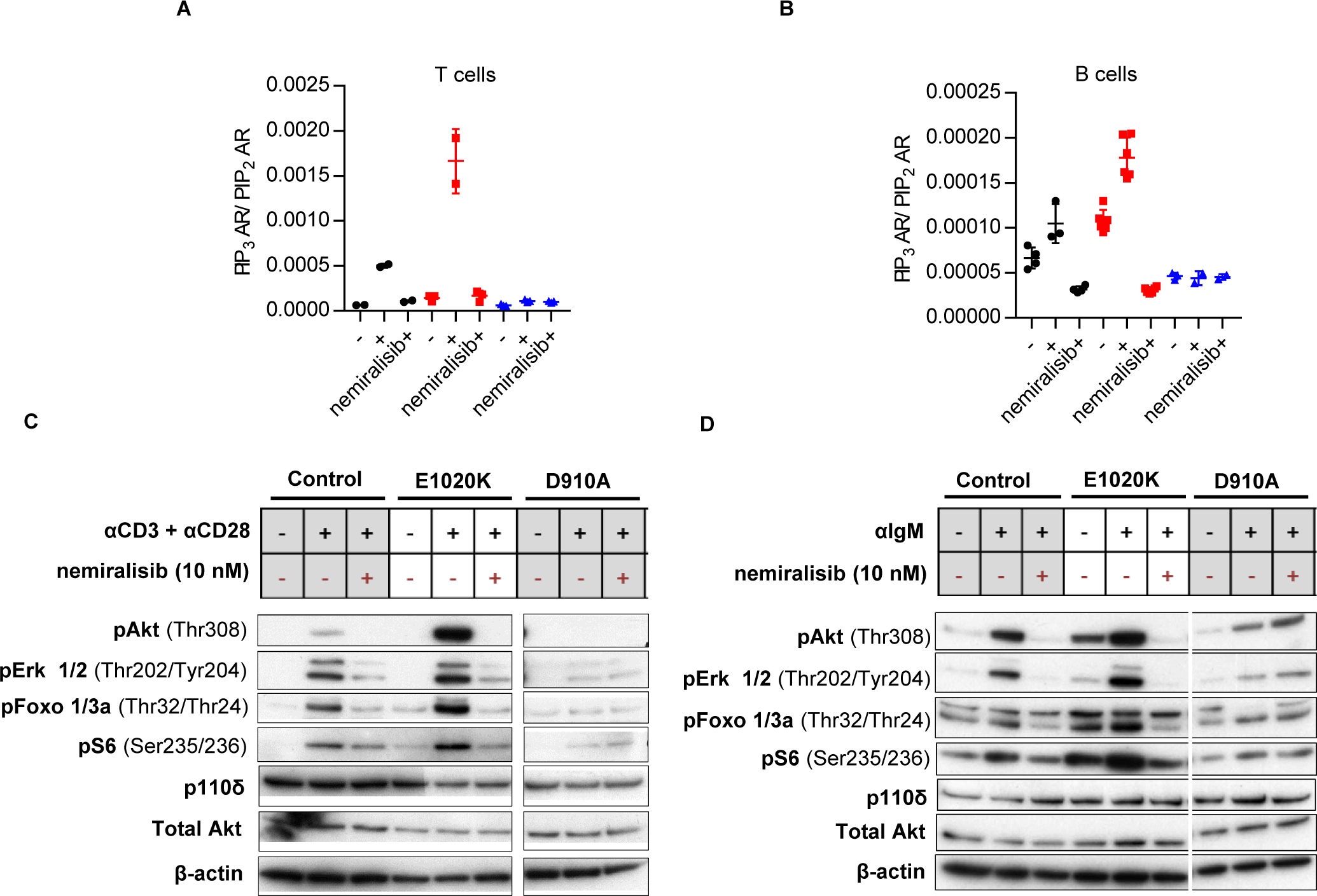
PI3Kδ hyper-activation leads to increased PIP_3_ and pAKT levels that can be reduced using a selective PI3Kδ inhibitor. **A:** PIP_3_ levels in purified T cells from wild-type, p110δ^E1020K^ and p110δ^D910A^ mice, unstimulated or stimulated with anti-CD3 and anti-CD28 in the presence or absence of the selective PI3Kδ inhibitor, nemiralisib (n=2-3). **B:** PIP_3_ levels in purified B cells from wild-type, p110δ^E1020K^ and p110δ^D910A^ mice unstimulated, or stimulated with anti-IgM in the presence or absence of nemiralisib (n=2-6). **C:** Western blots of purified T cells from wild-type, p110δ^E1020K^ and p110δ^D910A^ mice, unstimulated or stimulated with anti-CD3 and anti-CD28 in the presence or absence of nemiralisib. **D:** Western blots of purified B cells from wild-type, p110δ^E1020K^ and p110δ^D910A^ mice, unstimulated or stimulated with anti-IgM in the presence or absence of nemiralisib. (Representative of 2 independent experiments).

PIP_3_ binds to the protein kinase AKT, supporting its phosphorylation on Thr308 and subsequent activation. Western blotting of purified p110δ^E1020K-GL^ T cells showed increased AKT phosphorylation following stimulation with anti-CD3 and anti-CD28 antibodies compared to wild-type cells, whereas AKT phosphorylation in p110δ^D910A^ T cells was below the limit of detection (Fig 2C). In B cells, both basal and anti-IgM-induced phosphorylation of AKT were elevated in p110δ^E1020K-GL^ cells, while strongly diminished in p110δ^D910A^ B cells (Fig 2D). The phosphorylation of ERK and the AKT effector proteins, FOXO and S6, were similarly affected. All phosphorylation events in wild-type and p110δ^E1020K-GL^ cells were reduced to the levels observed in p110δ^D910A^ cells by inhibition with nemiralisib. As expected, p110δ protein expression was not affected by the E1020K or D910A mutations (Figs 2C, D).

Germline p110δ^E1020K-GL^ mice had near normal numbers of myeloid cells in the bone marrow and spleen. In the bone marrow we observed a significant B cell lymphopenia that was associated with a block in B cell development between Pro-B and Pre-B cells and did not extend to the spleen. Although these mice had normal numbers of splenic B cells, there was an increased proportion of marginal zone B and B1 cells with an altered distribution of transitional B cells (Supplementary Fig. 2). The thymus of p110δ^E1020K-GL^ mice was normal except for a mild reduction in single positive CD8^+^ T cells. In the spleens and lymph nodes there were increased proportions of activated/memory T cells identified by high CD44 expression and low CD62L expression, and increased numbers of Foxp3^+^ T regulatory cells (Supplementary Fig 3). These T cell and B cell phenotypes were recapitulated in p110δ^E1020K-T^ and p110δ^E1020K-B^ mice, respectively (Supplementary Figs 4 and 5.). The reciprocal effects of the inhibitory D910A and activating E1020K mutations on specific cell subsets demonstrate the pivotal role of PIP_3_ during lymphocyte development and highlight the importance of a tight control of PI3Kδ activity.

Analysis of serum immunoglobulins showed that p110δ^E1020K-GL^ mice had elevated levels of IgG1 and IgG2b and a trend towards increased levels of IgG2c, IgA, and IgE isotypes compared to wild-type mice. There was also a trend to hyper IgM in p110δ^E1020K-GL^ mice as is frequently observed in APDS patients ^10, 11, 14^, whereas p110δ^D910A^ mice were antibody deficient (Supplementary Fig 6). The level of serum IgG3, which has been shown to be protective against *S. pneumoniae^19^,* was comparable in p110δ^E1020K-GL^ and wild-type mice (Supplementary Fig 6).

### Susceptibility to *S. pneumoniae* is caused by PI3Kδ hyper-activation in B cells

Given that the immunological phenotype of p110δ^E1020K-GL^ mice strongly resembled that of APDS, we sought to determine whether these mice recapitulate the increased susceptibility to *S. pneumoniae* observed in APDS patients. We infected p110δ^E1020K-GL^, p110δ^D910A^, and p110δ^WT^ mice with *S. pneumoniae* (TIGR4 serotype 4) and followed their survival for 10 days (Fig 3A). Interestingly, despite being antibody-deficient, p110δ^D910A^ mice did not show increased susceptibility to *S. pneumoniae*. By contrast, p110δ^E1020K-GL^ mice showed accelerated disease onset and increased mortality (Fig 3A).

**Figure 3:**
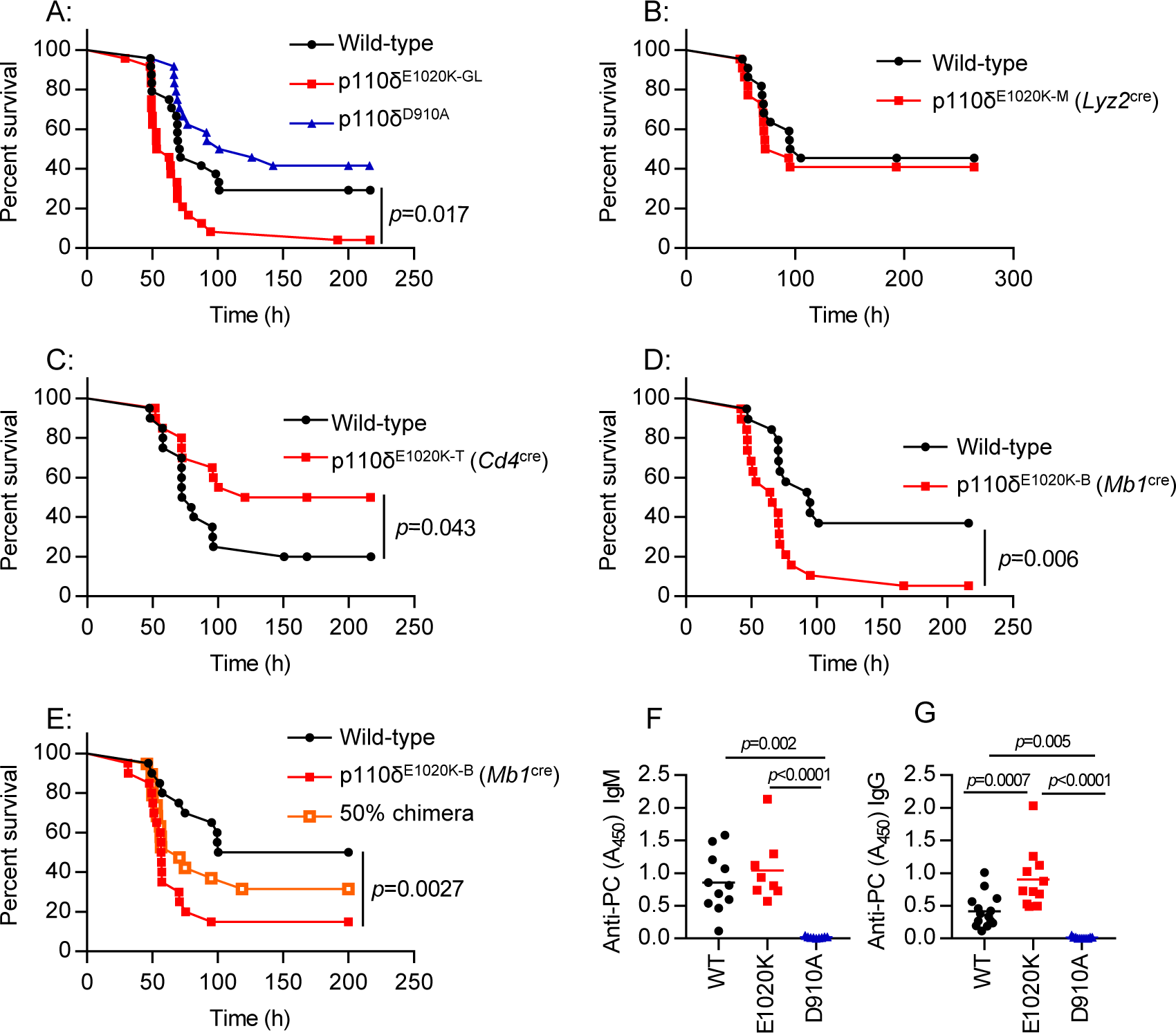
PI3Kδ hyper-activation leads to increased susceptibility to *S. pneumoniae* infection. **A**: Germline p110δ^E1020K-GL^ mice show accelerated disease development and significantly increased mortality compared to control mice in response *S. pneumoniae* infection, while kinase dead p110δ^D910A^ mice do not respond differently to wild-type mice. **B-D**: The p110δ^E1020K^ mutation was introduced conditionally into myeloid cells, T cells and B cells by crossing onto *Lyz2*^cre^, *Cd4*^cre^ and *Mb1*^cre^ lines respectively. The myeloid conditional mutation did not affect survival following *S. pneumoniae* infection in p110δ^E1020K-M^ mice (B), while T cell conditional p110δ hyper-activation lead to improved survival in p110δ^E1020K-T^ mice (C). However, introducing the p110δ^E1020K^ mutation specifically in B cells (p110δ^E1020K-B^ mice) replicated the increased susceptibility to *S. pneumoniae* seen in p110δ^E1020K-GL^ mice (D). **E**: Transfer of p110δ^E1020K-B^ bone marrow into irradiated RAG2^−/−^ recipients also conferred increased susceptibility to *S. pneumoniae* infection compared to recipients receiving wild-type bone marrow, and this phenotype was not fully rescued in a 50% BM chimera. **F-G:** Naïve PI3Kδ^E1020K^ mice produce normal levels of anti-phosphorylcholine IgM and significantly higher levels of anti-PC IgG, while PI3Kδ^D910A^ mice produce no natural antibody. (Data from 2 independent experiments combined. A: n=24; B: n=22; C: n=20; D: n=20; E: n=20; F-G: WT n=11; E1020K n=8 D901A n=9).

In order to determine the cell type responsible for increased susceptibility to *S. pneumoniae*, we next infected the lineage-restricted p110δ^E1020K-B^, p110δ^E1020K-T^ and p110δ^E1020K-M^ mice. Myeloid expression of p110δ^E1020K^ had no effect on the course of *S. pneumoniae* infection (Fig 3B), whereas expression of p110δ^E1020K^ in T cells was protective (Fig 3C). Only the p110δ^E1020K-B^ mice replicated the increased susceptibility of the p110δ^E1020K-GL^ mice to *S. pneumoniae* (Fig 3D). Furthermore, transfer of p110δ^E1020K-B^ bone marrow into irradiated RAG2^−/−^ recipients also conferred increased susceptibility to infection, which was only partially rescued by co-transferring wild-type and p110δ^E1020K-B^ bone marrow at a 1:1 ratio (Fig 3E). These results indicate that B cells drive the increased susceptibility to *S. pneumoniae* infection in p110δ^E1020K-GL^ mice in an immune-dominant manner.

Natural antibodies against phosphorylcholine (PC) can offer protection against infection with encapsulated bacteria, including *S. pneumoniae ^20, 21^*. We found that p110δ^D910A^ mice lacked anti-PC antibodies, presumably because of the absence of B1 and MZ B cells which are the major source of natural antibodies^19, 22–24^. By contrast, anti-PC antibodies in the serum from p110δ^E1020K^ mice were similar (IgM) or elevated (IgG)(Fig 3 F, G) compared to wild-type mice. Therefore, susceptibility to *S. pneumoniae* in p110δ^E1020K^ mice cannot be explained by a failure to produce natural antibodies against conserved bacterial epitopes.

### B cells drive early pathology during *S. pneumoniae* infection in an antibody-independent manner but are required to prevent chronic infection

In order to further investigate antibody-mediated protection in the context of PI3Kδ hyper-activation, we immunized mice with Pneumovax, a 23-valent polysaccharide vaccine^25^. Following infection with *S. pneumoniae*, wild-type mice were completely protected by this vaccination protocol, while p110δ^E1020K-GL^ mice were only partially protected, to a level similar to that in non-immunized wild-type mice. By contrast, p110δ^D910A^ mice did not benefit from vaccination (Fig 4A, B). Interestingly, p110δ^E1020K-GL^ mice and wild-type mice produced a similar antibody response, in contrast to p110δ^D910A^ mice that showed no response (Fig 4C). These data indicate that Pneumovax vaccination clearly protects against *S. pneumoniae*, as shown in immunized wild-type mice. Despite this protection p110δ^E1020K^ mice remained significantly more susceptible to *S. pneumoniae* infection. Together, these data suggest that B cells can affect susceptibility to *S. pneumoniae* by a mechanism that is at least in part antibody-independent.

**Figure 4:**
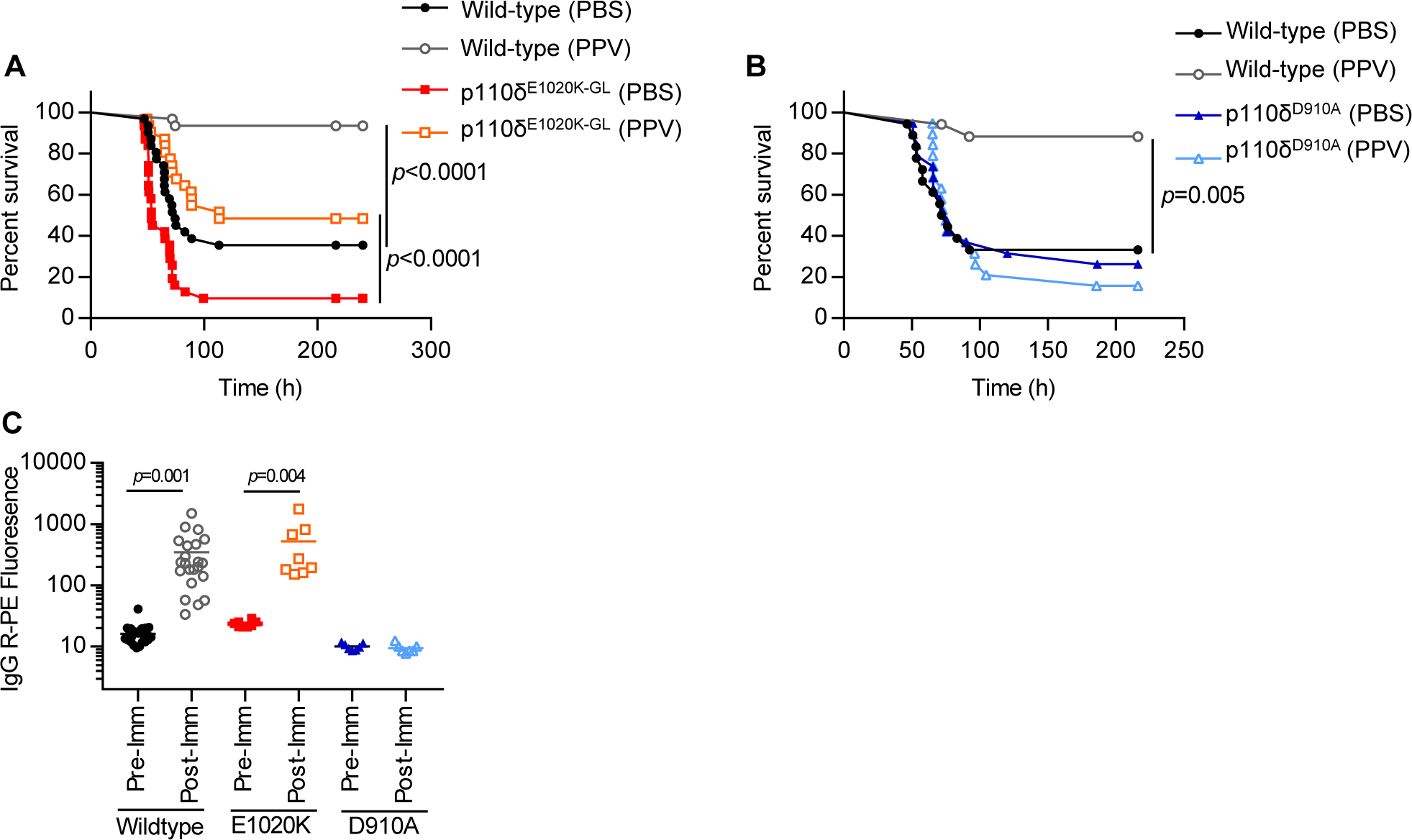
PI3Kδ^E1020K-GL^ mice respond to Pneumovax and have normal antibody levels. **A-B:** Pneumovax (T-independent pneumococcal polysaccharide vaccine, PPV) partially protects p110δ^E1020K-GL^ mice against infection (A), while p110δ^D910A^ are not protected (B). **C:** p110δ^E1020K-GL^ mice produce normal levels of total IgG in response to Pneumovax immunisation (anti-pneumococcal serotype 4 shown) while p110δ^D910A^ mice do not respond to vaccine. (A: results from 3 independent studies combined, n=30; B: results from 2 independent studies combined, n=19; C: results from 2 independent studies combined: WT n=22; E1020K n=8; D910A n=6)

To determine more definitively whether B cells can be pathogenic in the context of *S. pneumoniae* infection, we infected wild-type and *Ighm*^tm1^ (µMT) mice which lack mature B cells^26^. Strikingly, *Ighm*^tm1^ mice showed reduced susceptibility to *S. pneumoniae* infection, delaying disease onset from ∼2 days in wild-type mice to ∼5 days in *Ighm*^tm1^ mice (Fig 5). Although survival in *Ighm*^tm1^ mice was increased up to 30 days post infection compared to wild-type mice, CFU counts from the lungs of mice surviving to this time-point indicates that, despite appearing clinically healthy, 41% (7/17) of *Ighm*^tm1^ mice failed to clear the infection compared to 100% clearance in wild-type mice (Fig 5). Taken together, these results suggest that during early time-points in the local infected environment B cells can be pathogenic, while at later stages they are required to prevent chronic infection.

**Figure 5:**
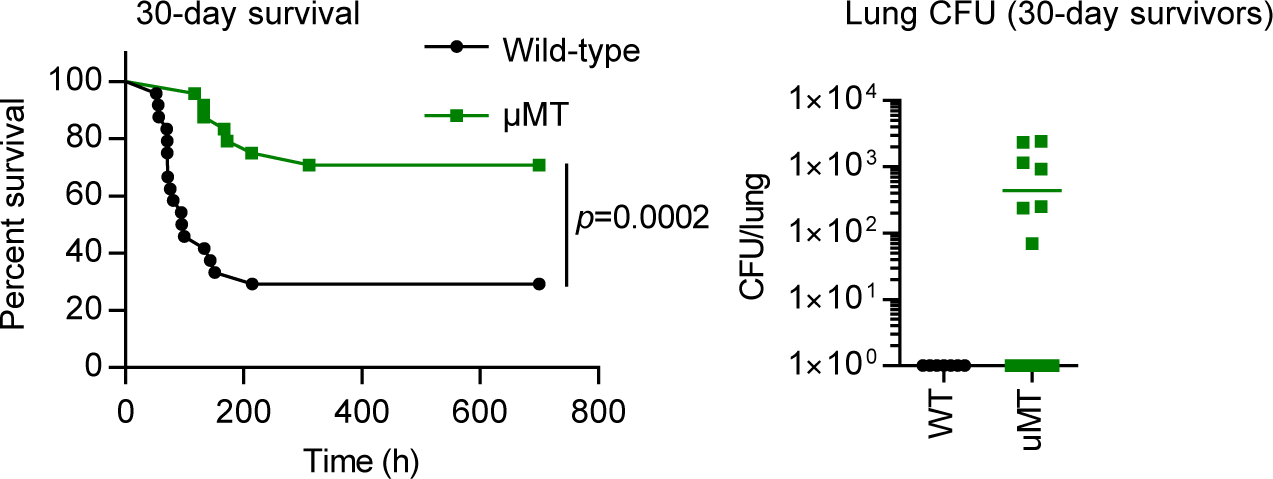
B cells could drive increased pathology in response to *S. pneumoniae* infection, independent of bacterial clearance. *Ighm*^tm1^ (µMT) mice show delayed disease progression and improved overall survival up to 30 days post *S. pneumoniae* infection, however 41% (7/17) surviving *Ighm*^tm1^ mice failed to clear bacteria from the lung tissue at 30 days post infection compared to 100% clearance in surviving wild-type mice. (Data from 1 study, n=24)

Increased susceptibility to *S. pneumoniae* could either be due to uncontrolled bacterial proliferation or be caused by an aberrant immune response to the pathogen. The bacterial titers from lungs and spleens of wild-type, p110δ^D910A^ and p110δ^E1020K-GL^ mice were similar at 24h post infection, suggesting that the different susceptibilities did not correlate with different abilities to control bacterial outgrowth during the early phase of infection (Supplementary Fig 7A). We observed a trend towards increased levels of TNFα, IL-6, IL-1β and IL-1α in the lung tissue of p110δ^E1020K-GL^ mice at 24h post infection (Supplementary Fig 7B). Consistent with this, the levels of TNFα, IL-6 and IL-1β were reduced in the lungs of wild-type mice treated with nemiralisib prior to *S. pneumoniae* infection (Supplementary Fig 7C), indicating that, while PI3Kδ signaling affect the amount of pro-inflammatory cytokines produced in response to infection, the increase in response to PI3Kδ hyper-activation is modest.

### CD19^+^ B220^−^ B cells produce high levels of IL-10 in response to *S. pneumoniae* infection in comparison with conventional CD19^+^ B220^+^ B cells

We hypothesized that increased susceptibility to *S. pneumoniae* infection can be mediated by a specific subpopulation of B cells. Therefore, we studied the B cell compartment in various tissues and found an atypical population of CD19^+^B220^−^ B cells that was rare in wild-type mice, but significantly increased in p110δ^E1020K-GL^ mice and was absent in p110δ^D910A^ mice (Fig 6A). In the spleen and bone marrow, there was a 10- and 5-fold increase, respectively, in the numbers of CD19^+^B220^−^ B cells in p110δ^E1020K-GL^ mice compared to wild-type mice (Fig 6A). Infection with *S. pneumoniae* did not induce further expansion of CD19^+^B220^−^ B cells in the lungs or other tissues examined 24h post-infection (Fig 6B).

**Figure 6:**
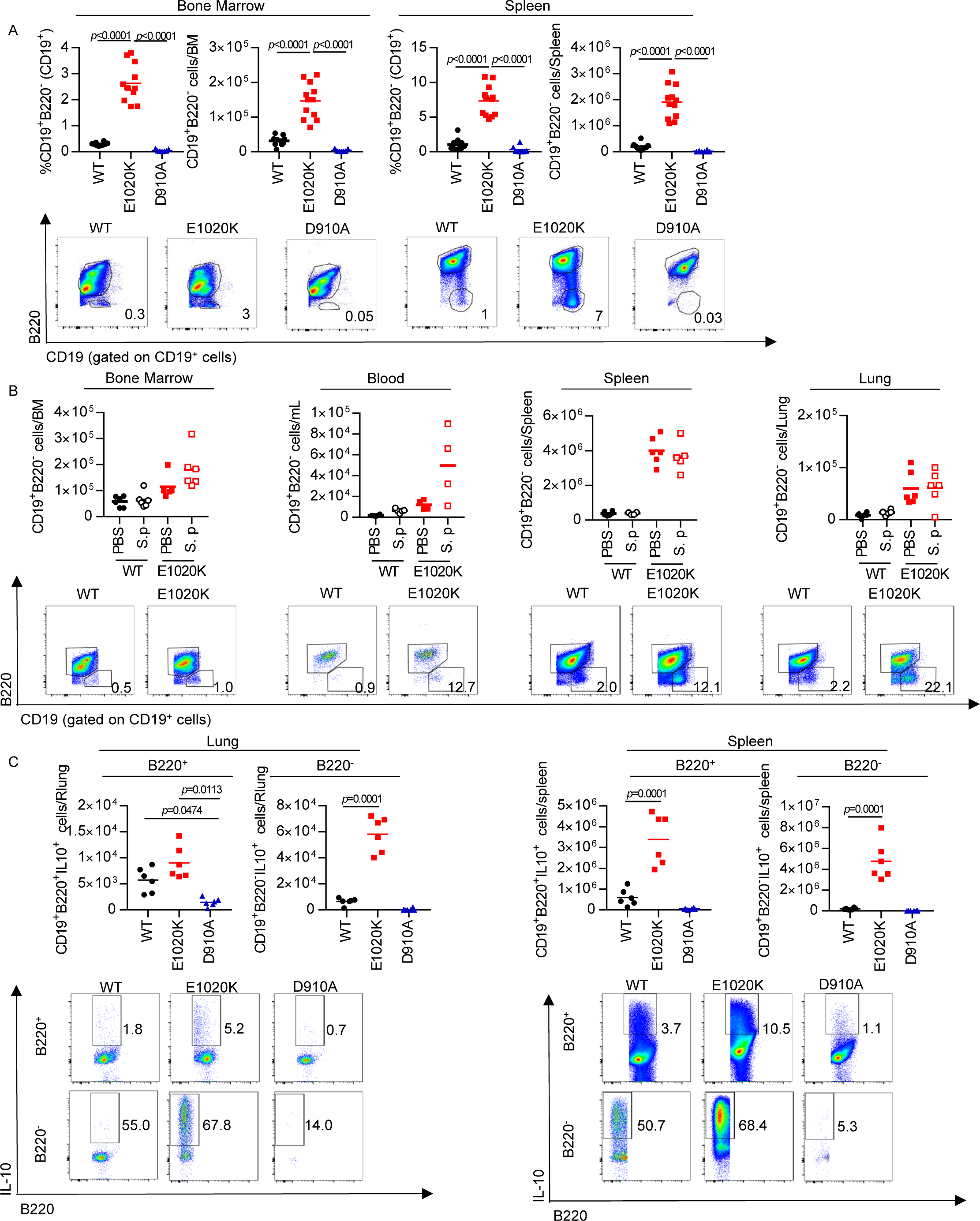
p110δ^E1020K^ mice have an expanded population of IL-10 producing CD19^+^B220^−^ B cells. **A:** Naïve p110δ^E1020K^ mice show a significant increase in the proportion and number of B220^−^ B cells in bone marrow and spleen. While this population is detectable in wild-type mice, it is absent from p110δ^D910A^ mice. Representative pseudocolor plots show mean cell proportions in naïve mice **B:** *S. pneumoniae* (S.p) infection does not lead to significant expansion or recruitment of B220^−^ B cells at 24h post infection in bone marrow, blood, spleen or lung tissue. Representative pseudocolor plots show mean cell proportions in naïve mice. **C:** At 24h post *S. pneumoniae* infection, IL-10 production is increased in p110δ^E1020K^ B cells from both lung and spleen, and reduced in p110δ^D910A^ B cells compared to wild-type mice. Furthermore, B220^−^ B cells produce higher levels of IL-10 in response to S. *pneumoniae* infection compared to B220^+^ B cells in wild-type and p110δ^E1020K^ mice. Representative pseudocolor plots show mean cell proportions in *S. pneumoniae* infected mice (A: combined data from 2 independent experiments, WT n=10; E1020K n=12; D910A n=7, B: representative data from 2 independent experiments, n=6, C: representative data from 2 independent experiments n=6).

To ascertain if these cells were also present in the lungs and to distinguish resident from circulating cells, we labelled circulating leukocytes in wild-type and p110δ^E1020K-GL^ mice by intravenous injection of biotin-conjugated anti-CD45. We then stained the lung homogenate with fluorochrome-conjugated anti-mouse CD45 and streptavidin, and used flow cytometry to distinguish tissue resident leukocytes from those present in the lung capillaries. We found a significant increase in the proportion and number of tissue-resident, but not circulating, B cells in p110δ^E1020K-GL^ mice compared to wild-type mice. There was an increase in both CD19^+^B220^+^ and CD19^+^B220^−^ cells among the tissue-resident B cells which demonstrates that both populations can take residence in the lung (Supplementary Fig 8).

Given that B cells from p110δ^E1020K-GL^ mice drive susceptibility to *S. pneumoniae* in an antibody-independent manner, we sought to look at other properties of B cells. We found a trend towards lower IL-10 protein levels and a significant 10-fold decrease in IL-10 mRNA expression at 24h post infection in whole lung homogenates from wild-type mice treated with nemiralisib (Supplementary Fig 7C, D), indicating that PI3Kδ signaling can regulate IL-10 levels in the lung post infection. IL-10 is an important immune-regulatory cytokine known to affect the course of *S. pneumoniae* infection^27, 28^ and therefore increased IL-10 production could explain the B cell-dependent but antibody-independent effects seen in p110δ^E1020K-GL^ mice. To investigate this further, we crossed the p110δ^E1020K-GL^ and p110δ^D910A^ mice with a highly sensitive IL-10 reporter (*Il10*^ITIB^) mouse^29^.

At 24h post *S. pneumoniae* infection, the frequencies of the IL-10-producing B cells were significantly increased in the lungs of p110δ^E1020K-GL^ *Il10*^ITIB^ mice compared to in wild-type *Il10*^ITIB^ mice (Supplementary Fig 7E). In addition, IL-10-producing B cells were barely detected in lungs from p110δ^D910A^ mice (Supplementary Fig 7E). Frequencies of the IL-10-producing CD11b^+^ myeloid, T and NK cells in lungs were variable, but not consistently increased in p110δ^E1020K-GL^ mice (Supplementary Fig 7E). Further analysis of the CD19^+^ B cell subset isolated from lungs and spleens at 24h post *S. pneumoniae* infection showed that the proportion of IL-10-producing cells was increased among the B220^−^ B cell subset as compared to the B220^+^ B cell subset in p110δ^E1020K-GL^ mice (5% B220^+^ vs 68% B220^−^) and wild-type mice (2% B220^+^ vs 55% B220^−^) (Fig 6C). Furthermore, the proportion and absolute number of IL-10-producing CD19^+^B220^−^ cells and CD19^+^B220^+^ cells was increased in p110δ^E1020K-GL^ mice compared to wild-type mice, while such IL-10-producing cells were virtually absent in p110δ^D910A^ mice (Fig 6C). This indicates that CD19^+^B220^−^ cells are the predominant population of B cells that produce IL-10 in a PI3Kδ-dependent manner.

Previously, B cells that produce IL-10 have been termed B regulatory cells (Bregs)^30^. We sought to compare the phenotype of CD19^+^B220^−^ IL-10-producing B cells with conventional IL-10-producing CD19^+^B220^+^ Bregs described previously (Fig 7). Following *in vitro* stimulation for 5h and analysis of cell surface markers and intracellular IL-10 we found that there were clear phenotypic differences between conventional B220^+^ Breg cells and the B220^−^ B cells we describe here, including the differential expression of CD43 and IgM (Fig 7). The lack of B220 expression and low IgM expression differentiate the CD19^+^B220^−^ IL-10-producing B cells from conventional B1 cells^31^.

**Figure 7:**
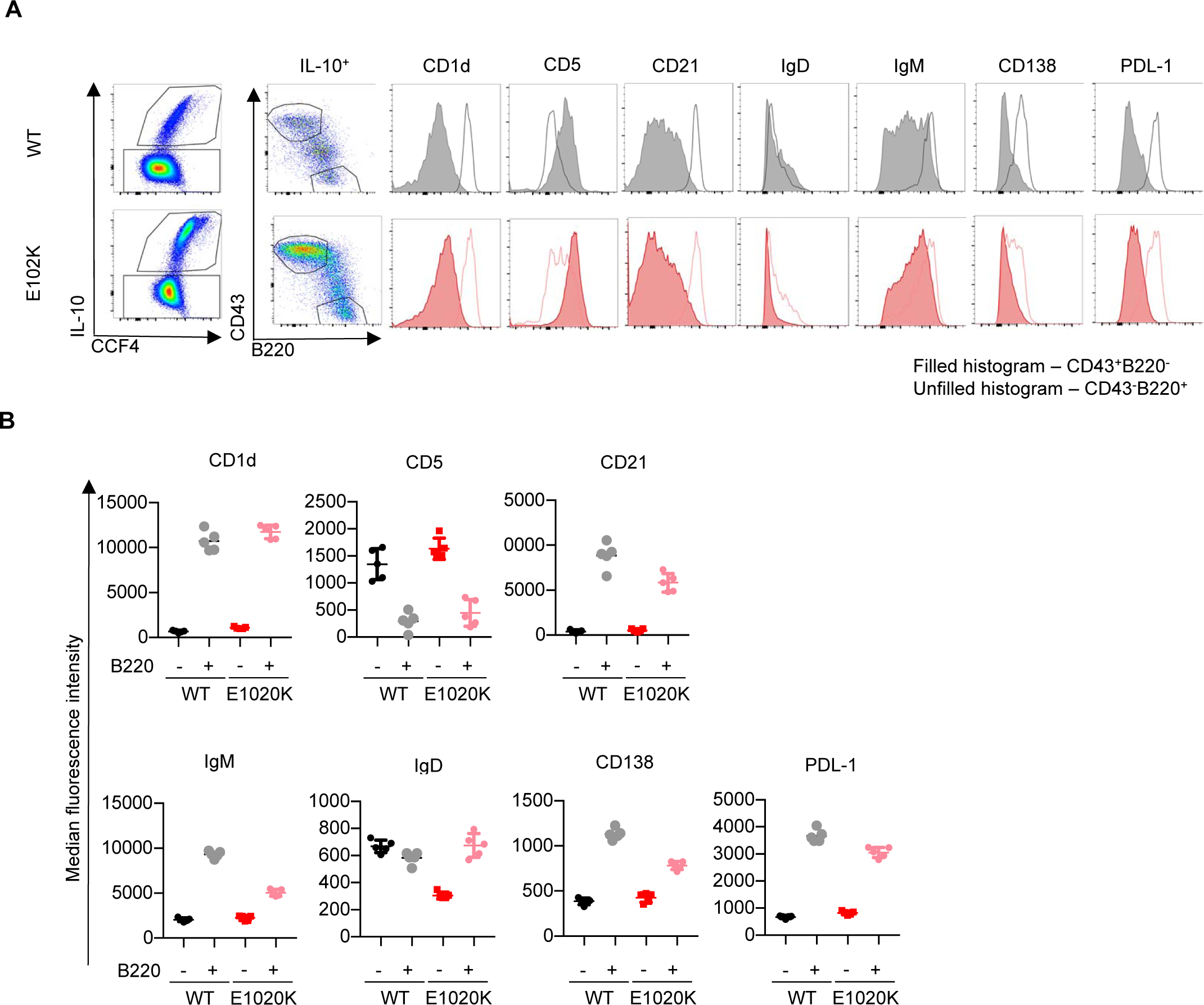
IL-10 producing B220^−^ B cell subset is a novel B cell subset, expanded in PI3Kδ^E1020K^ mice. In order to compare surface marker expression from B220^−^ and B220^+^ IL-10 producing B cells, splenocytes from *Il10*^ITIB^ mice (WT and E1020K) were stimulated with LPS/PdBu/Ionomycin/Brefeldin A for 5 hours and then stained for cell surface markers as shown. **A:** After gating on IL-10 producing B cells, CD43^++^B220^−^ cells were compared to CD43^−^B220^+^ cells. Histograms highlighting the differential surface marker expression as measured by median fluorescence intensity (MFI) between these populations are shown. **B:** Comparison of surface marker MFI show that CD19^+^ B220^−^ IL-10^+^ B cells express: CD43^++^CD5^int/+^ CD23^−^CD21^−^CD1d^lo/int^ IgM^+/−^IgD^lo/−^PDL1^−^CD138^−^ as opposed to conventional Bregs expressing: CD19^hi^ B220^hi^ IL-10^+^ CD43^−^ CD5^Var^CD23^−^CD21^++^CD1d^hi^ IgM^hi^ IgD^Var/lo^ PDL1^+^CD138^int^. (Data from 1 experiment, n=5)

### Nemiralisib ameliorates the susceptibility to *S. pneumoniae* in p110δ^E1020K^ mice

Given that increased PI3Kδ activity in mice leads to increased mortality after *S. pneumoniae* infection we explored if nemiralisib would provide protection. Treatment of p110δ^E1020K-GL^ mice with inhaled nemiralisib 24h prior to infection led to a 20% increase in survival. While nemiralisib treatment did not affect the numbers of CD19^+^B220^−^ B cells, the proportion of IL-10 producing CD19^+^B220^−^ B cells in the lungs were reduced in comparison to non-treated mice (Fig 8A). This was in keeping with our prior observations that nemiralisib treatment reduced IL-10 mRNA levels in the lung tissue of *S. pneumoniae* infected wild-type mice (Supplementary Fig 7D), suggesting a role for IL-10-producing CD19^+^B220^−^ B cells in the PI3Kδ-dependent susceptibility to infection.

**Figure 8:**
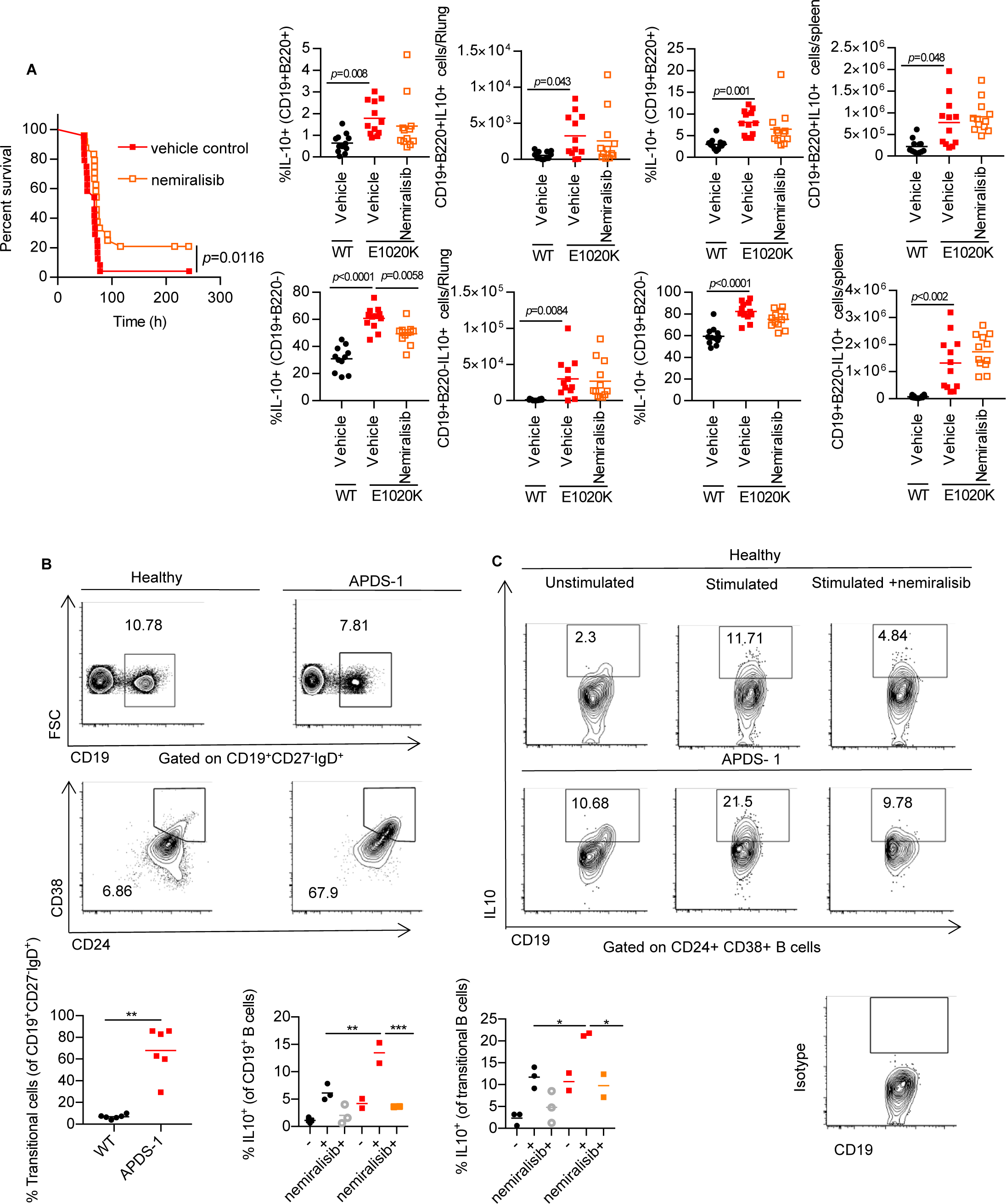
Nemiralisib treatment reduces the proportion of IL-10 producing B cells in mice and humans. **A:** Prophylactic treatment with nemiralisib improved survival in p110δ^E1020K-GL^ mice and was associated with a significant reduction in the proportion of IL-10 producing B220^−^ B cells in the lung at 24h post infection. (A: Combined data from two independent studies, survival n=24; lung tissue analysis n=12) **B:** Blood from patients with APDS (n=6) and healthy controls (n=6) was obtained and the B cell phenotype was determined by flow cytometry. Transitional B cells are identified as CD19^+^IgD^+^CD27^−^CD24^+^CD38^+^. Representative contour plots with outliers show the mean proportions of cells. **C**: Freshly isolated PBMCs were unstimulated or stimulated with plate bound anti-CD3 and anti-IL-2 for 72h in the presence or absence of nemiralisib (n=2-3). The proportion of IL-10 producing cells among the total B cell and transitional B cell populations was determined by flow cytometry. Representative contour plots with outliers show mean proportions of IL-10 producing cells. (Combined data from 2 independent experiments).

### Nemiralisib treatment suppresses IL-10 production in B cells of APDS patients

Cohort studies have shown that 75% of patients with APDS have elevated circulating transitional B cells (CD19^+^IgM^++^CD38^++^)^14^. Other studies have shown that such transitional B cells can produce high levels of IL-10^32, 33^. We obtained blood samples from patients with APDS and healthy controls and confirmed that all of these APDS patients had elevated proportions of transitional B cells (Fig 8B). We isolated PBMCs from two of these APDS patients and three healthy controls and stimulated them for 72h with anti-CD3 and IL-2 ^32^. After stimulation we found more IL-10-producing B cells and more IL-10-producing transitional B cells in PBMCs of APDS patients than in PBMCs of healthy controls, while treatment with nemiralisib effectively suppressed IL-10 production in these cells (Fig 8C). Therefore, in keeping with reduction of CD19^+^B220^−^IL-10 producing B cells in the mice, nemiralisib can also reduce IL-10 producing human transitional B cells.

## Discussion

Our results show that enhanced PI3Kδ signaling leads to the increased susceptibility to *S. pneumoniae* infection through a B cell-dependent, but antibody-independent mechanism. We found an expansion of an aberrant B cell population which could be equivalent to the elevated transitional B cell population found in APDS patients. These cell populations may have a detrimental effect during the early phase of *S. pneumoniae* infection.

### B cells drive increased susceptibility to *S. pneumoniae* infection in p110δ^E1020K^ mice

In this study, we show that B cells can increase the susceptibility to *S. pneumoniae* infection during the first few hours after exposure. Mice were more susceptible to *S. pneumoniae* infection when the hyperactive p110δ^E1020K^ mutation was expressed in B cells. By contrast, mice expressing the p110δ^E1020K^ mutation only in T cells were protected. The basis for the protection remains unknown and is the focus of ongoing study. The p110δ^E1020K^ mice have increased numbers of T_FH_ and Treg cells and the conventional T cells have a more activated phenotype. A recent study has documented a population of lung-resident innate Th17 cells that confers protection against re-challenge with *S. pneumoniae*^34^. Whether such a subset is affected by PI3Kδ activity and can also protect against immediate challenge is not known. However, we did not detect increased proportions of Th17 cells after infection. Regardless of the mechanism, the protection offered by T cells was overcome by the adverse effects of B cells in p110δ^E1020K-GL^ mice.

We describe a subset of B cells which lack the common B cell marker B220, and whose abundance is correlated with the susceptibility to *S. pneumoniae* infection. Moreover, we show that the development of this CD19^+^B220^−^ B cell subset is highly dependent on the level of activity of PI3Kδ. Susceptibility to *S. pneumoniae* infection was not altered in p110δ^D910A^ mice as compared to wild-type mice. Kinase-dead p110δ^D910A^ mice lack natural and anti-capsular antibodies which may make them more susceptible, however, their lack of CD19^+^B220^−^ B cells may counterbalance this antibody deficiency, such that overall the mutation has little net effect on *S. pneumoniae* susceptibility.

### PI3Kδ signaling controls IL-10 production in B cells

A large proportion of the CD19^+^B220^−^ B cells produced IL-10. Although increased IL-10 could potentially reduce the production of inflammatory cytokines, we found similar or increased levels of TNFα, IL-6 and IL-1 in the lungs of p110δ^E1020K^ mice and no difference in CFU counts at 24h post infection.

One of the defining characteristics of APDS patients is an increased proportion of CD24^+^CD38^+^ B cells defined as transitional B cells ^10–12, 14, 15^. Intriguingly, production of IL-10 is a characteristic of these B cells as well. High IL-10 levels in response to secondary *S. pneumoniae* infection following influenza A is associated with increased lethality when compared to a primary *S. pneumoniae* infection, and importantly, the outcome of a secondary *S. pneumoniae* infection is improved by IL-10 neutralisation^27, 28^. This indicates that, while IL-10 is required for normal immune regulation and resolution of inflammation^35^, excess IL-10 during the early stages of *S. pneumoniae* infection could have an acute detrimental effect. This fits with our observation that nemiralisib treatment improved the outcome of mice infected with *S. pneumoniae* and reduced IL-10 mRNA levels in the lung. Further studies are required to investigate the time and location specific effects of B cell dependent IL-10 production, keeping in mind that B cells have the potential to produce a number of different cytokines (such as IL-6, TNF-α, and IL-35) that could affect *S. pneumoniae* susceptibility^36^.

We postulate that the transitional B cells which are expanded in APDS patients may not just be precursors to more mature B cells, but also include cells that are functionally equivalent to the CD19^+^B220^−^ cells that we have identified in p110δ^E1020K^ mice and thus may also contribute to the increased susceptibility to *S. pneumoniae* infection and hence to the high incidence of bronchiectasis characteristic of this disease^15^. In this context, it is interesting that CD19^+^B220^−^ B cells had previously been described as immature progenitors in the bone marrow^37^, rather than a functional PI3Kδ dependent subset found in peripheral tissues as we describe here. Although the CD19^+^B220^−^ B cells we describe resemble B1 cells by some criteria, such as the expression of CD5 and CD43, their lower IgM expression, lack of B220 and distribution pattern suggest that they are related but distinct to conventional B1 and B10 cells^30, 31^. A similar subset has previously been described as being dependent on CD19 and increased in the absence of PTEN expression and PTEN haploinsuffiency in humans can lead to an APDS-like syndrome^38–40^.

Our results do not question the important role that antibodies play in the protection against *S. pneumoniae* and other encapsulated bacteria. Indeed, in wild-type mice, immunization with Pneumovax offered complete protection. Rather, our results highlight a hitherto underappreciated subset of B cells that increase the pathological response shortly after infection. It remains to be determined whether this is a feature of the B cells being lung-resident (and hence among the first cells of the immune system to encounter the invading pathogens) and by which mechanisms B cells either condition lung epithelia to become more susceptible to invasion of *S. pneumoniae*, reduce the clearance of *S. pneumoniae* by phagocytes and/or increase the collateral adverse effects of the early innate immune response to this pathogen. This is in keeping with the increasing realization that B cells are important sources of different cytokines and can modulate immune responses independently of their capacity to present antigen and produce antibodies^36^.

### Therapeutic implications

Intriguingly, the inhaled inhibitor nemiralisib offered protection in not just in p110δ^E1020K^ but also in wild-type mice infected with *S. pneumoniae*, when administered prior to infection. This result suggests that preventive inhibition of PI3Kδ can be beneficial not only just in the context of APDS, but also in other conditions associated with PI3Kδ hyperactivation, such as COPD^41^.

The results presented herein are relevant to our understanding and the potential treatment of patients with APDS. Clinical trials are currently ongoing using both inhaled and systemic PI3Kδ inhibitors (NCT02593539, NCT02435173 and NCT02859727). Systemic use of PI3Kδ inhibitor has shown promise in reduction of lymphoproliferation and immune cell aberrations in patients with APDS, however, no comments were made regarding recurrent infections or respiratory manifestations^42^. Systemic use of PI3Kδ inhibitors in malignancies have been associated with serious side effects, including colitis^43^. It would therefore be of importance to determine whether inhaled PI3Kδ inhibitors could alleviate the respiratory manifestations of APDS while reducing the risk of the adverse effects associated with systemic inhibition. Moreover, the results may also be relevant to other conditions, such as COPD^17^, where PI3Kδ may be activated by non-genetic mechanisms. Overall, our findings suggest that in subjects with increased PI3Kδ activity prophylactic administration of PI3Kδ inhibitors may help alleviate the course of *S. pneumoniae* infection.

## Materials and Methods

### Mouse strains and gene targeting

The p110δ^E1020K^ mice were generated by Ozgene, Australia using homologous recombination in ES cells. A duplicate sequence corresponding to the last coding exon in *Pik3cd* and carrying the E1020K mutation was flanked by loxP sites and inserted 3’ to the wild-type sequence. Upon Cre-mediated recombination, the wild-type sequence is replaced by the mutant E1020K sequence. p110δ^D910A^, *Cd4*^cre^, *Mb1*^cre^, *Lyz2*^cre^, *Ighm*^tm1^ (µMT) and *Il10*^ITIB^ mice have been described previously^19, 26, 29, 44–46^. Throughout the study, the p110δ^E1020K^ allele was heterozygous whereas the p110δ^D910A^ alleles were homozygous. Genotyping was performed by Transnetyx (Cordova, TN).

### *S. pneumoniae* stock preparation

*Streptococcus pneumoniae* (TIGR4, serotype 4 (provided by Professor Jeremy Brown, University College, London)) was grown to mid-log phase (OD_500_ = 0.5-0.7) in Todd-Hewitt broth (Oxoid) supplemented with 0.5% yeast extract (Oxoid) at 37°C, 5% CO_2_. The bacteria were collected by centrifugation, and resuspended in PBS/20% glycerol (Sigma-Aldrich) prior to freezing in liquid N_2_ and storage at -80 °C. Stocks were assessed for viable CFU counts and homogeneity by plating out serial dilutions of three frozen samples on blood agar plates (LB agar, supplemented with 5% defibrinated sheep blood (Oxoid)) after incubation for 24h at 37 °C, 5% CO_2_. *S. pneumoniae* colonies were confirmed by the presence of an α-hemolytic zone and sensitivity to optochin (Sigma-Aldrich). Virulent stocks were maintained by performing an *in vivo* passage every 6-12 months.

### S. pneumoniae infections

Frozen stocks were thawed and washed twice by centrifugation in sterile PBS before resuspending at 4×10^7^ CFU/mL in PBS. The suspension was kept on ice at all times, and used for infection within 2h of thawing (no loss of viability was observed under these conditions). Mice were lightly anaesthetized by inhalation of 3% isoflurane and maintained with 2% isoflurane. Mice (males and females aged 8-12 weeks) were infected intranasally with 50µL *S. pneumoniae* suspension containing 2×10^6^ CFU. Animals were observed to confirm inhalation of the dose and full recovery from anesthesia. The infection dose was routinely confirmed by plating out serial dilutions of the inoculum on blood agar plates, as described above.

### Survival studies

Mice were infected with *S. pneumoniae* as described above. Pre-infection body weights were recorded and animals were weighed daily post infection. Animals were monitored three times a day for a period of 10 days post infection. Disease progression was assessed by assigning clinical scores without knowledge of the individual genotypes: 0: Healthy; 1: mild clinical signs; 2: Up to 2 moderate clinical signs; 3: up to 3 moderate signs. Animals were culled when they showed >25% bodyweight loss or reach score 3. The most frequently observed clinical signs were piloerection, hunched posture, tremor, and labored breathing. At the study end-point, animals were culled and terminal blood samples collected.

### Bone Marrow transfer

A single cell suspension of p110δ^E1020K-B^ and wild-type (C57Bl/6.SJL) donor bone marrow was prepared in sterile HBSS (Sigma-Aldrich) as described below (Isolation of immune cells from mouse tissues). Cells from two sex-matched donors were pooled. RAG2^−/−^ recipient mice were supplied with 4mg/mL Neomycin (Sigma-Aldrich) in drinking water before sub-lethal irradiation (one dose of 500Rads over 63 seconds), and received 3×10^6^ donor cells by intravenous (tail vein) injection. Neomycin treatment was maintained for 4 weeks post transfer, and after 8 weeks reconstitution was confirmed by analyzing tail bleeds.

### Vaccination

For vaccine studies, mice were immunized with Pneumovax II (pneumococcal polysaccharide vaccine, serotypes: 1-5, 6B, 7F, 8, 9N, 9V, 10A, 11A, 12F, 14, 15B. 17F, 18C, 19F, 19A, 20, 22F, 23F and 33F; Sanofi Pasteur MSD). One 0.5mL dose containing 50µg/mL of each serotype was diluted 1:12.5 in sterile PBS to 4µg/mL. Mice were given one 100µL (0.4µg) dose by intraperitoneal injection, 14 days before infection with *S. pneumoniae.* Blood samples were taken from the tail vein prior to vaccination and 14 days after vaccination into serum collection tubes (BD microtainer, SST) samples were centrifuged at 15000xg for 5 min and the serum stored at -20°C until analysis.

### Nemiralisib treatment

The PI3Kδ inhibitor nemiralisib (GSK2269557) was supplied by GSK, Respiratory Refractory Inflammation DPU, UK. The nemiralisib suspension was prepared on the day of use in 0.2% Tween80, and administered intranasally to mice twice daily at 0.2mg/kg in a total volume of 50µL, as described above (*S. pneumoniae* infections). Unless otherwise specified, the first dose was given 24h before infection, and treatment was maintained for the duration of the study.

### Isolation of immune cells from mouse tissues

Mice were euthanized by CO_2_ inhalation and cervical dislocation. Lungs were perfused with 10mL cold PBS through the right ventricle, and collected into cold PBS. Lungs were homogenized using a GentleMACS tissue dissociator and mouse lung dissociation enzyme kit from Miltenyi according to the manufacter’s instructions. The homogenate was transferred to 15mL tubes (BD Falcon) and washed by centrifugation (500xg) in 10mL cold PBS. The pellet was resuspended in 3mL 37.5% isotonic Percoll (Sigma) at room temperature and centrifuged at 650xg for 20min with low acceleration and no brake. The supernatant including tissue debris was removed, and the cell pellet was washed and resuspended in cold PBS. Where CFU counts were required, the right lung was processed as described above and the left lung was processed for CFU counts as described below. Bone marrow cells were collected by flushing cold PBS through femurs and tibias collected, filtered through 40µm cell strainers and washed by centrifugation in 5mL PBS. Spleens, thymus and peripheral lymph nodes were homogenized in PBS by pushing the tissue through 40µm cell strainers (BD) using a syringe plunge. The cell suspension was then transferred to a 15ml Tube (BD falcon) and washed once in 5ml cold PBS. For blood, bone marrow and spleen, red blood cells (RBC) were lysed using hypotonic ammonium chloride RBC lysis buffer (Sigma). Lysing was quenched with cold PBS and the cells were collected by centrifugation. Single cell suspensions were processed for flow cytometry as described below.

### S. pneumoniae CFU counts

Lungs were homogenized in 1ml PBS using a Bullet Blender using 3x 3mm steel beads: speed 8, 3min (Next Advance, USA). Spleens were homogenized as described above in 2mL PBS. Serial dilutions (10-fold) were performed for spleen and lung homogenates, and samples were plated out on blood agar as described above. Plates were incubated for 24h at 37°C and *S. pneumoniae* colonies were counted.

### *In vitro* stimulation of mouse B cells

Splenocytes were isolated from naïve wild-type, p110δ^E1020K-GL^ and p110δ^D910A^ mice that had been crossed with the *Il10*^ITIB^ reporter line as described above and the total cell count was obtained using a CASY counter. The cells were resuspended in complete RPMI (RPMI plus 5% (v/v) FCS, 50 μM β-mercaptoethanol and 100 μg/ml penicillin and streptomycin) and plated at 5×10^6^ cells in 100µL per well in a 96-well U bottom plate. The cells were stimulated with 10 ng/ml LPS, 50ng/ml PdBu (Sigma, USA), 0.25 μg/ml Ionomycin (Sigma, USA) and 1 μl/ml of Brefeldin A (eBioscience) for 5h at 37 °C. The cells were then processed for IL-10 detection and flow cytometry as described below.

### Detection of IL-10 in *Il10*^ITIB^ reporter mice

Single cell suspensions from *S. pneumoniae* infected mice or *in vitro* stimulated cells were prepared as described above. The cells were incubated with 3.3µM CCF4-AM, a Fluorescence Resonance Energy Transfer substrate for β-lactamase (LiveBLAzer kit, Invitrogen), and 3.6mM probenecid (Sigma) in complete RPMI for 90min at 29°C, as previously described ^29^, then placed on ice and collected by centrifugation. Cells were then processed for flow cytometry as described below.

### Flow cytometry

Antibodies used for flow cytometry are listed in Table 1. Single cell suspensions were stained using an antibody master mix in PBS/0.5%BSA for 40min at 4°C. Cells were washed and fixed in 4% paraformaldehyde (Biolegend) for 10min at room temperature before washing 2x in PBS/0.5%BSA. For intracellular detection of IL-10 in human samples, the cells were fixed and permeabilized with Cytofix/Cytoperm buffer (BD Biocsiences, USA) following surface staining. For FoxP3 staining, the eBioscence FoxP3/Transcription Factor staining Buffer Set was used according to the manufacturer’s instructions. Non-fluorescent counting beads (AccuCount Blank Particles 5.3µm; Spherotech, USA) were added to quantify absolute cells numbers. Samples were kept at 4°C until analysis (BD Fortessa5). Analysis was carried out using FlowJo (Treestar) analysis software.

**Table 1:** Antibodies.

| **anti-mouse** | **Clone** | **Supplier** |
| --- | --- | --- |
| CCR2 | 475301 | R&D Systems |
| CD11b | M1/70 | Biolegend |
| CD11c | N418 | Biolegend |
| CD138 | 281-2 | Biolegend |
| CD19 | ID3 | BD Biosciences |
| CD1d | K253 | Biolegend |
| CD21 | 7E9 | Biolegend |
| CD23 | B3B4 | Biolegend |
| CD25 | PC61 | Biolegend |
| CD28 | 37.51 | Biolegend |
| CD3 | 2C11 | Biolegend |
| CD4 | GK1.5 | Biolegend |
| CD43 | S7 | BD Biosciences |
| CD44 | IM7 | Biolegend |
| CD45 | 30-F11 | Biolegend |
| CD45R(B220) | RA3-6B2 | BD Biosciences |
| CD5 | 53-7-3 | eBioscience<br>(ThermoFisher) |
| CD62L | MEL-14 | Biolegend |
| CD8a | 53-6.7 | Biolegend |
| FoxP3 | FJK-16s | eBioscience<br>(ThermoFisher) |
| IgD | 11-26c.2a | Biolegend |
| IgM | II/41 | eBioscience<br>(ThermoFisher) |
| IL-7R | A7R34 | Biolegend |
| KLRG1 | 2F1 | Biolegend |
| Ly6C | HK1.4 | Biolegend |
| Ly6G | 1A8 | Biolegend |
| Ly6G/Ly6C(Gr1) | RB6-8C5 | Biolegend |
| NK1.1 | PK136 | Biolegend |
| Siglec-F | E50-2440 | BD Biosciences |
| TCRβ | H57-597 | Biolegend |
| $\gamma\delta$TCR | UC7-13D5 | Biolegend |
| **anti-human** | **Clone** | **Supplier** |
| CD19 | SJ25C1 | BD Biosciences |
| CD24 | ML5 | BD Biosciences |
| CD38 | HIT2 | BD Biosciences |
| IL-10 | JES3-9D7 | Miltenyi Biotec |

### *In vitro* stimulation of human B cells

Freshly isolated PBMCs from patients and control individuals were stimulated with 0.5μg/ml purified plate bound anti CD3 monoclonal antibody (clone-OKT3) (Invitrogen) and 20ng/ml recombinant human IL2 in presence or absence of, the p110δ inhibitor, nemiralisib (10nM) for 72h. This stimulation led to the activation of CD40L (CD154) on CD4 T cells. The interaction between CD40L on CD4 T cells with CD40 on B cells resulted in the production of IL-10 in regulatory B cells. For a final stimulation 50ng/ml PdBu (sigma, USA), 0.25 μg/ml ionomycin (Sigma, USA) and 1μl/ml of Brefeldin A (eBioscience) were also added for the last 4h. Cells were washed, surface stained, fixed and permeabilized with Cytofix/Cytoperm buffer (BD Biocsiences, USA) for the intracellular detection of IL-10 as described above.

### Natural Antibody ELISA

Anti-phosphorylcholine IgM and IgG levels were assessed in serum from naïve mice. Blood samples were collected by cardiac puncture and serum collected as described above. NUNC Maxisorp ELISA plates were coated overnight at 4°C with 20µg/mL phosphorylcholine-BSA (BioSearch) in sodium bicarbonate coating buffer (pH9) (Biolegend) (100µL/well). Plates were washed 4x with PBS/0.2% Tween20 followed by blocking for 1h at room temperature in PBS/1%BSA. Serum samples were diluted 1:10 in PBS/1% BSA and pooled WT serum was diluted from 1:5 to 1:400 to confirm the linear range of the assay. After removing the blocking solution, samples were added at 100µL/well, and the plates were incubated overnight at 4°C. The plates were then washed 4x in PBS/0.2%Tween20. Polyclonal HRP conjugated goat-anti-mouse IgM (abcam ab97023) or IgG (abcam ab97230) was diluted 1:5000 in PBS/1% BSA, added at 100µL per well and the plates were incubated for 2h at room temperature. The plates were washed 6x with PBS/0.2%Tween20, TMB substrate (Biolegend) was added at 100µL per well and incubated for 5-10 min at room temperature. The reaction was stopped by adding 50µL 2N H_2_SO_4_ solution. Absorbance was read at 450nm (Fluostar omega, BMG Labtech).

### Tissue resident cell analysis (lung)

Mice received an intravenous (tail vein) injection of 3µg anti-mouse CD45 conjugated to biotin (Biolegend, clone 30-F11) in 100µL PBS. 4min after injection animals were killed and the lungs were collected into cold PBS without prior perfusion. Lungs were then finely minced using a scalpel blade and the pushed through a 100µm cell strainer (BD) the homogenate was collected in a 15mL tube (BD falcon) and washed in 10ml cold PBS. The pellet was resuspended in 3mL 37.5% isotonic Percoll (Sigma) at room temperature and processed as described above. The cells were stained for flow cytometry as described above, including streptavidin APC to identify anti-CD45-Biotin labelled circulating cells.

### Cytokine analysis

Cytokine levels were measured in the supernatant from lung homogenates prepared for CFU analysis. The homogenates were centrifuged for 1min at 10000xg to remove tissue debris. Samples collected from wild-type mice treated with nemiralisib were analyzed using Meso Scale Discovery mouse Th1/Th2 9-plex ultrasensitive kits, according to manufacturer’s instructions. Samples collected from genetically modified mice were analyzed using the flow cytometry based Legendplex mouse inflammation 13-plex panel (Biolegend).

### Gene Array

Wild-type mice were treated with nemiralisib and infected with *S. pneumoniae* as described above. At 24h post infection, whole lungs were collected and snap frozen in liquid nitrogen. Samples were randomized and analyzed (Affymetrix Genechip Mouse genome 430 2.0 Array) by Expression Analysis, Quintiles Global Laboratories. Data analysis was performed by Computational Biology and Statistics, Target Sciences, GlaxoSmithKline, Stevenage. Data was normalised using the robust multiarray average method^47^ and quality checked in R/Bioconductor^48^ using the *affy* package^49^. A linear model was fitted to the RMA normalised data for each probset and differiential expression analysis was conducted in the ArrayStudio software (Omicsoft Corporation). P-values were false discovery rate corrected by the method of Benjamini and Hochberg^50^. Probes with an absolute fold change >1.5 and an adjusted p-value <0.05 were called significant. The data been deposited in the National Center for Biotechnology Information Gene Expression Omnibus (GEO) and are accessible through GEO series accession number GSE109941.

### Luminex antibody analysis

Measurement of IgG recognizing pneumococcal polysaccharide of 13 serotypes was performed as described previously ^51^. The assay was modified in order to measure murine IgG as follows: 50µl of 1:10 diluted mouse sera was added to each well containing the pneumococcal polysaccharides coupled to microspheres and the secondary antibody used was 50µl of 10µg/ml goat F(ab’)_2_ anti-mouse IgG (H+L)- R-phycoerythrin (Leinco Technologies, Inc). R-PE fluorescence levels were recorded for each serotype and median fluorescence intensities were compared for the different genotypes.

### PIP_3_ quantification

CD4 and CD8 cells were isolated from mouse splenocytes using immunomagnetic negative selection by biotinylated cocktail of antibodies and Streptavidin dynabeads (Invitrogen). T cells were stimulated with anti-CD3 (1mg/ml), (145-2c11, Biolegend) anti- CD28 (2µg/ml), (35.51, Biolegend) antibodies followed by crosslinking with anti- Armenian hamster IgG (10 µg/ml, Jackson ImmunoResearch labs). Cells were stimulated for 1 min at 37°C. B cells were similarly isolated from mouse splenocytes and stimulated for 1 min at 37 °C with anti- IgM 2µg/ml (AffinPure F(ab’)_2_ Fragment goat anti-mouse IgM from Jackson Immuno-Research labs).

After stimulation of the cells, we terminated the reactions by addition of 750µl kill solution (CHCl_3_:MeOH:1M HCL (10:20:1) and immediately froze the samples on dry ice. PIP_3_ levels were quantified by mass spectrometry as previously described^52^.

### Western blot

Purified T and B cells were isolated from murine spleens and stimulated as described above (PIP_3_ measurement). For performing western blots the stimulation period was 5 min. Isolated T cells were stimulated with anti -CD3 1µg/ml and anti -CD28 (2 µg/ml) with or without 10 Nm nemiralisb; or B cells with anti- IgM 4µg/ml (AffinPure F(ab’)_2_ Fragment goat anti-mouse IgM from Jackson ImmunosResearch labs).

Stimulated T and B cells were lysed with ice-cold lysis buffer (50mM HEPES, 150mM NaCl, 10mM NaF, 10mM Indoacetamide, 1% IGEPAL and proteinase inhibitors (Complete Ultra tablets, Roche)) for 15-20 min. Lysates were centrifuged at 15,000g for 10 min at 4 °C and supernatants were mixed with NuPage LDS sample buffer (life technologies). Samples were heated at 70°C and resolved on 4-12% NuPage bis-tris gel (Invitrogen), transferred to PVDF membranes and probed with the following antibodies: pAKT (T308, Cell Signaling, 1 in 1000 dilution); total AKT1 (2H10, Cell signaling, 1 in 2000 dilution); p110δ (Sc7176, Santa Cruz Biotechnology, 1 in 200 dilution); pS6 (S235/236, Cell Signaling, 1 in 500 dilution); pFoxo1/3a (T24/T32, Cell Signaling, 1 in 1000 dilution); pErk (p44/42, Cell Signaling, 1 in 200 dilution); βActin (Sc47778, Santa Cruz Biotechnology, 1 in 2000 dilution).

## Statistics

Data analysis was performed in Graphpad Prism. Where two groups were compared we used student’s t-test with Welch’s correction. Where three or more groups were compared we used 1-way ANOVA with Tukey’s multiple comparisons test. Survival data was analyzed using the Gehan-Breslow-Wilcoxon test. Statistical significance is indicated by asterisk as follows: *p* > 0.05 *; *p* ≤ 0.05 **; *p* ≤ 0.01 ***; *p* ≤ 0.001 ****

## Ethics

Animal experiments were performed according to the Animals (Scientific Procedures) Act 1986, licence PPL 70/7661.

Informed consent was obtained from patients and healthy controls. This study conformed to the Declaration of Helsinki and according local ethical review document 12/WA/0148.

## Acknowledgements

We thank Rahul Roychoudhuri, Bart Vanhaesebroeck and Martin Turner for their invaluable advice on the draft manuscript. We also thank Hicham Bouabe and Jürgen Heesemann (LMU Munich) for providing the *Il10*^ITIB^ mice and for advice. We thank Ramkumar Venigalla for advice on B cell phenotyping and Alice Denton for advice on the detection of tissue resident cells in the lung. Rainer Döffinger provided advice on the use of Luminex for the quantification of antibodies against pneumococcal serotypes; Keith Burling helped quantify mouse serum immunoglobulins. We gratefully acknowledge the support from the Babraham Institute Biological Services Unit, Biological Chemistry, Mass Spectroscopy and Flow Cytometry Facilities. Funding for the project was from the Medical Research Council MR/M012328/2 (AS, AMC, SN, KO), Wellcome Trust 103413/Z/13/Z and 206618/Z/17/Z (AC), 095691/Z/11/Z (KO), 095198/Z/10/Z (SN) Biotechnology and Biological Sciences Research Council BBS/E/B/000 -C0407, -C0409, -C0427 and -C0428 (KO). EBH is currently employed by Cambridge University Hospitals NHS Foundation Trust but is seconded to spend 50% of his time on GSK clinical trial research. He receives no other benefits or compensation from GSK. EBH was funded by the Wellcome Trust Translational Medicine and Therapeutics (TMAT) PhD program. SN is also supported by the National Institute for Health Research (NIHR) Cambridge Biomedical Research Centre. MRC is supported by the National Institute of Health Research (NIHR) Cambridge Biomedical Research Centre and the NIHR Blood and Transplant Research Unit and by a Medical Research Council New Investigator Research Grant (MR/N024907/1) and an Arthritis Research UK Cure Challenge Research Grant (21777). SS, GB, EBH, JNH and EMH are employed by GSK.

---

**Supplementary Figure 1:**
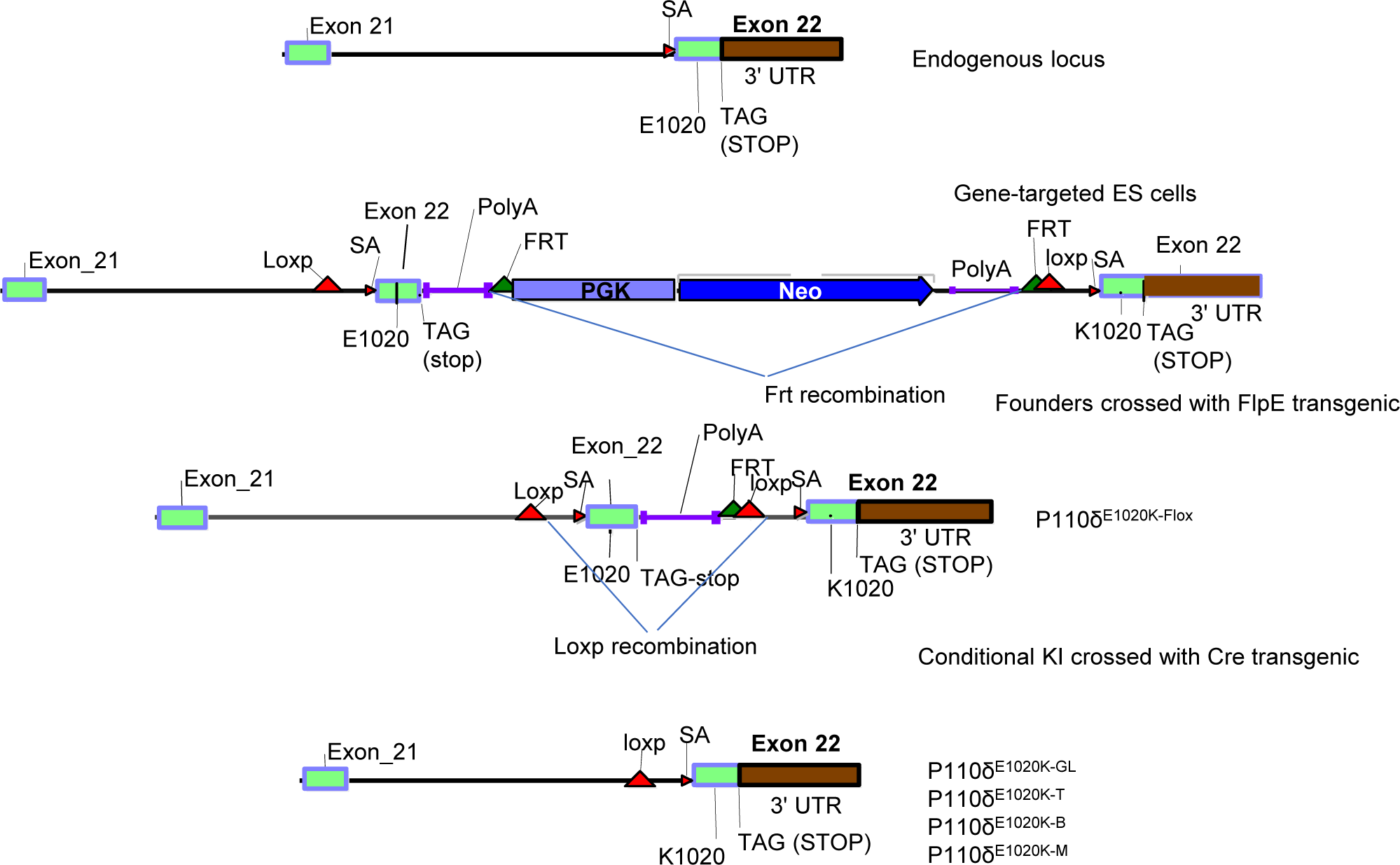
Gene targeting strategy for generating conditional p110δ^E1020K^ mice. The p110δ^E1020K^ mice were generated by OzGene using homologous recombination in ES cells. A duplicate sequence corresponding to the last coding exon in *Pik3cd* was flanked by loxP sites and inserted 3’ to the original sequence. The original sequence encoding E1020 was mutated to K1020. Upon Cre-mediated recombination, the wild-type sequence is replaced by the mutant E1020K sequence. In this study, we used *Tnfrsf4*^cre^ to delete in the germline, *Cd4*^cre^ to delete in T cells, *Mb1*^cre^ to delete in B cells and *Lyz2*^Cre^ to delete in myeloid cells.

**Supplementary figure 2:**
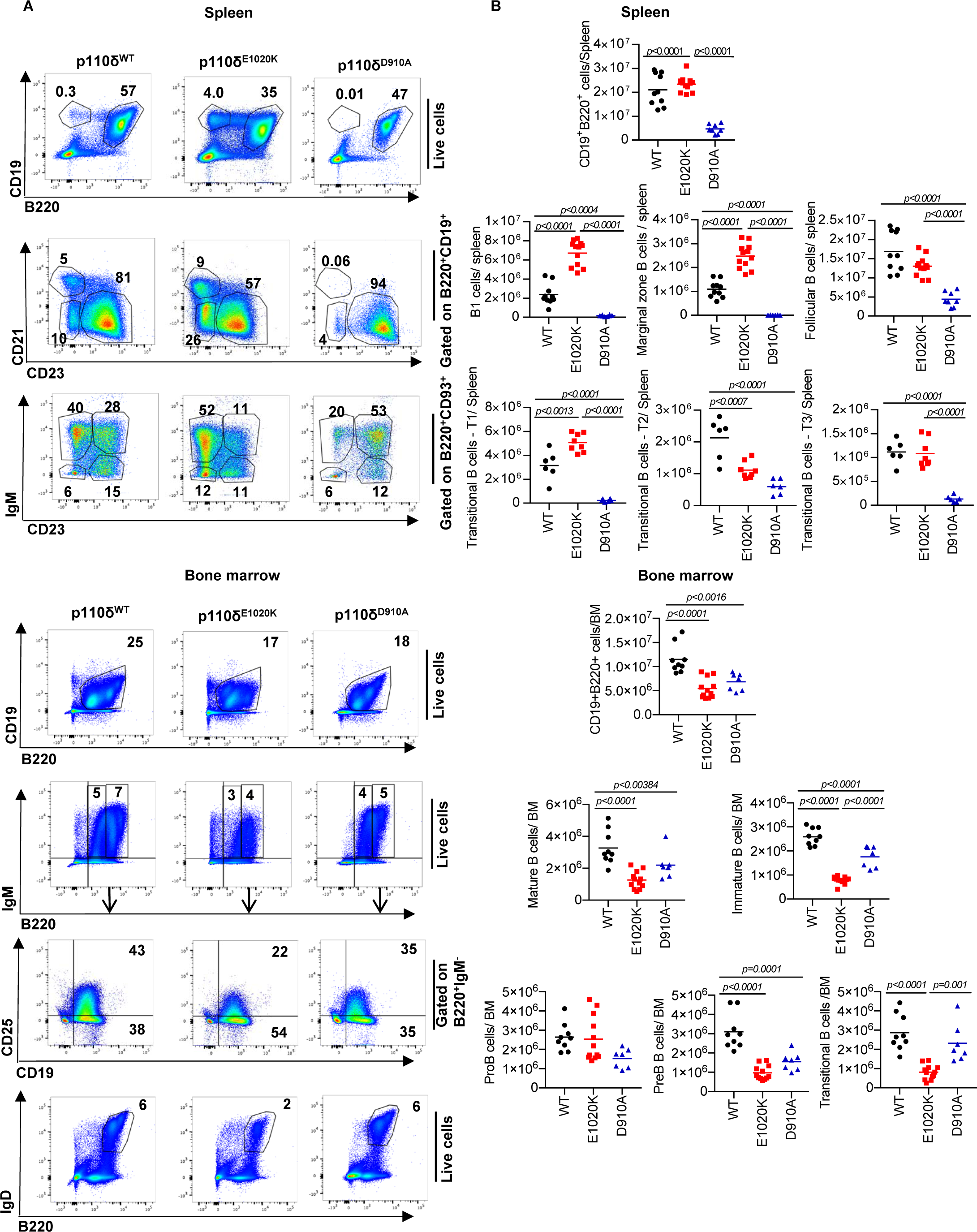
Phenotype of B cells in p110δ^E1020K-GL^ mice. **A:** B cell subsets in the spleen and bone marrow from wild-type, p110δ^E1020K-GL^ and p110δ^D910A^ mice were analyzed by flow cytometry and representative pseudocolor plots with the mean cell proportion are shown. **B:** In the spleen, total CD19^+^B220^+^ B cells were increased in p110δ^E1020K-GL^ mice, due to increased numbers of B1, marginal zone (MZ) and T1 transitional cells while follicular B cell numbers were normal. These populations were reduced in p110δ^D910A^ mice. Analysis of the bone marrow showed normal pro-B cell numbers in p110δ^E1020K-GL^ mice with a reduced number of pre-B cells, immature, transitional and mature B cells. Populations of cells are described as follows: Splenic B cells: Total B cells CD19^+^B220^+^, B1 cells CD19^+^B220^+^CD23^−^CD21^−^, Follicular B cells CD19^+^B220^+^CD23^+^CD21^+^, Marginal zone B cells CD19^+^B220^+^CD23^−^CD21^+^, Transitional T1 B cells B220^+^CD93^+^IgM^+^CD23^−^, Transitional T2 B cells B220^+^CD93^+^IgM^+^CD23^+^, Transitional T3 B cells B220^+^CD93^+^IgM^−^CD23^−^; Bone marrow B cells - Immature B cells CD19^+^B220^lo^IgM^+^, Mature B cells CD19^+^B220^hi^IgM^+^, Pro-B cells B220^+^IgM^−^CD19^+^CD25^−^, Pre-B cells B220^+^IgMCD19^+^CD25^+^, Transitional B cells B220^+^IgD^+^. (Combined data from 2 independent experiments, n=8-12).

**Supplementary figure 3:**
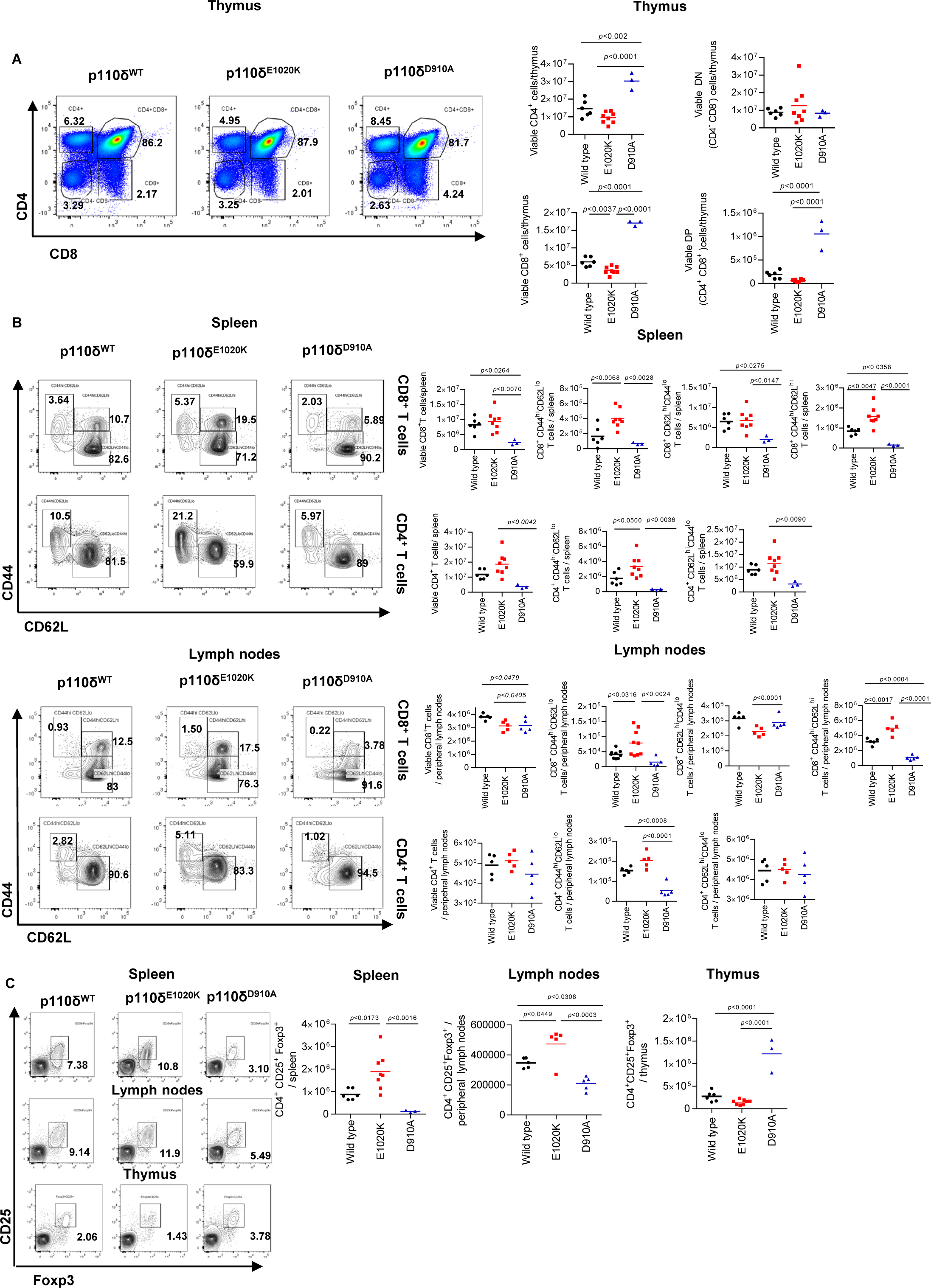
Phenotype of T cells in p110δ^E1020K-GL^ mice. T cell populations in the thymus, spleen and lymph nodes of wild-type, p110δ^E1020K-GL^ and p110δ^D910A^ mice were analyzed by flow cytometry. Representative pseudocolor or contour plots with outliers show mean cell proportions. **A:** Thymic T cell numbers in p110δ^E1020K^ mice were normal apart from a reduction in single positive CD8^+^ cells. **B:** The number of activated CD4^+^ and CD8^+^ T cells (CD44^hi^CD62^lo^) as well as memory CD8^+^ T cells (CD44^hi^CD62L^hi^) in lymph nodes and spleen were increased in p110δ^E1020K^ mice. **C:** The number of CD4^+^CD25^+^Foxp3^+^ regulatory T cells was increased in the lymph nodes and spleen of p110δ^E1020K^ mice. (Representative data from 2 independent experiments, n=3-8)

**Supplementary figure 4:**
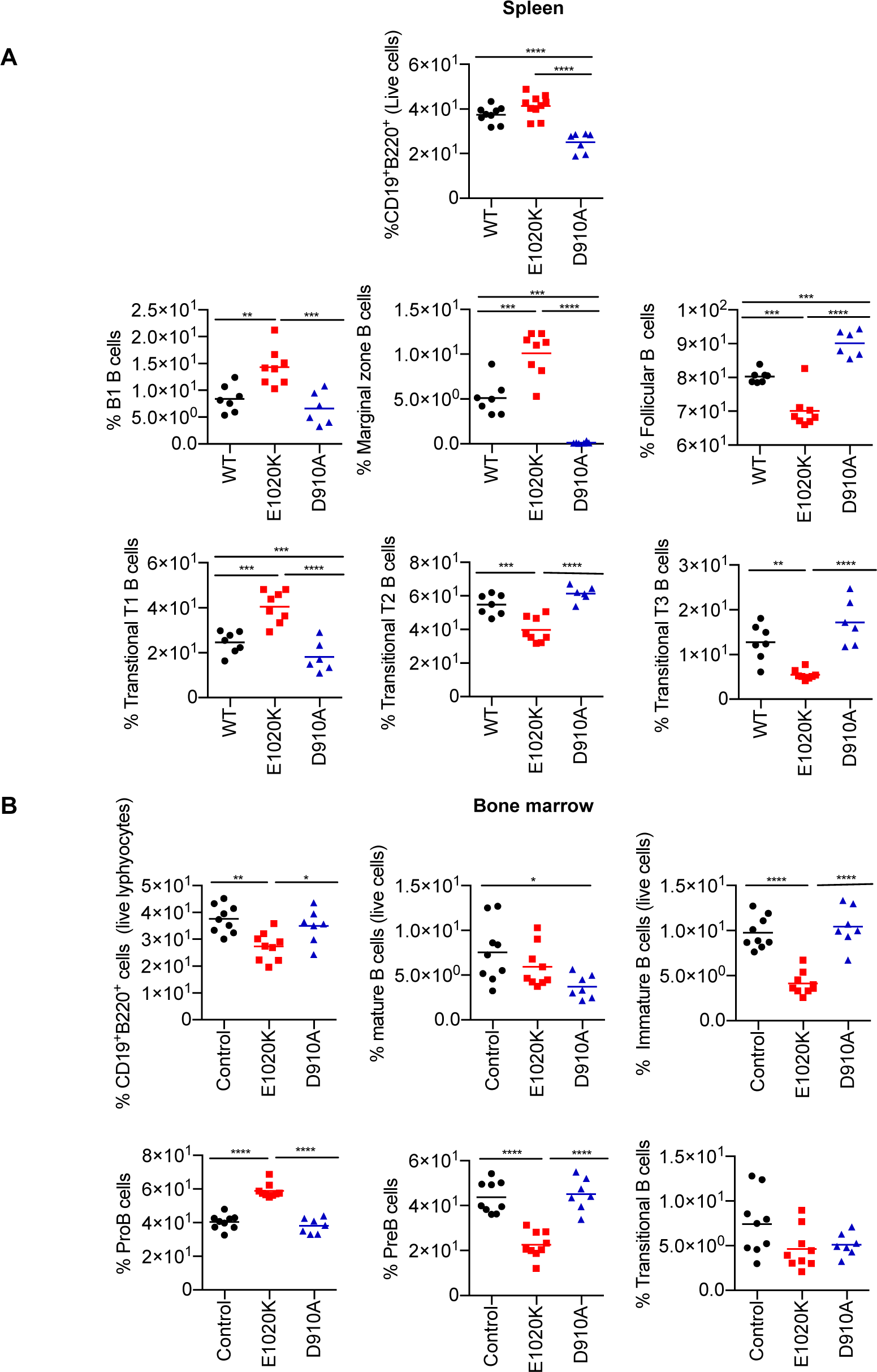
Phenotype of B cells in p110δ^E1020K-B^ mice. B cell subsets in the spleen and bone marrow from wild-type, p110δ^E1020K-B^ and p110δ^D910A^ mice were analyzed by flow cytometry, and was shown to recapitulate the phenotype of p110δ^E1020K-GL^ mice. **A:** In the spleen, the proportion of B1, marginal zone (MZ) and T1 transitional cells were increased, while the proportion of follicular B cells were reduced. **B:** Analysis of the bone marrow showed increased proportions of pro-B cells in p110δ^E1020K-B^ mice with reduced proportions of pre-B cells and immature B cells, and a trend towards reduced proportions of transitional and mature B cells. Populations of cells are described as follows: Splenic B cells - Total B cells CD19^+^B220^+^, B1 cells CD19^+^B220^+^CD23^−^CD21-, Follicular B cells CD19^+^B220^+^CD23^+^CD21^+^, Marginal zone B cells CD19^+^B220^+^CD23^−^CD21^+^, Transitional T1 B cells B220^+^CD93^+^IgM^+^CD23^−^, Transitional T2 B cells B220^+^CD93^+^IgM^+^CD23^+^, Transitional T3 B cells B220^+^CD93^+^IgM^−^CD23^−^; Bone marrow B cells - Immature B cells CD19^+^B220^lo^IgM^+^, Mature B cells CD19^+^B220^hi^IgM^+^, Pro-B cells B220^+^IgM^−^CD19^+^CD25^−^, Pre-B cells B220^+^IgMCD19^+^CD25^+^, Transitional B cells B220^+^IgD^+^. (Combined data from 2 independent experiments, n=6-10).

**Supplementary figure 5:**
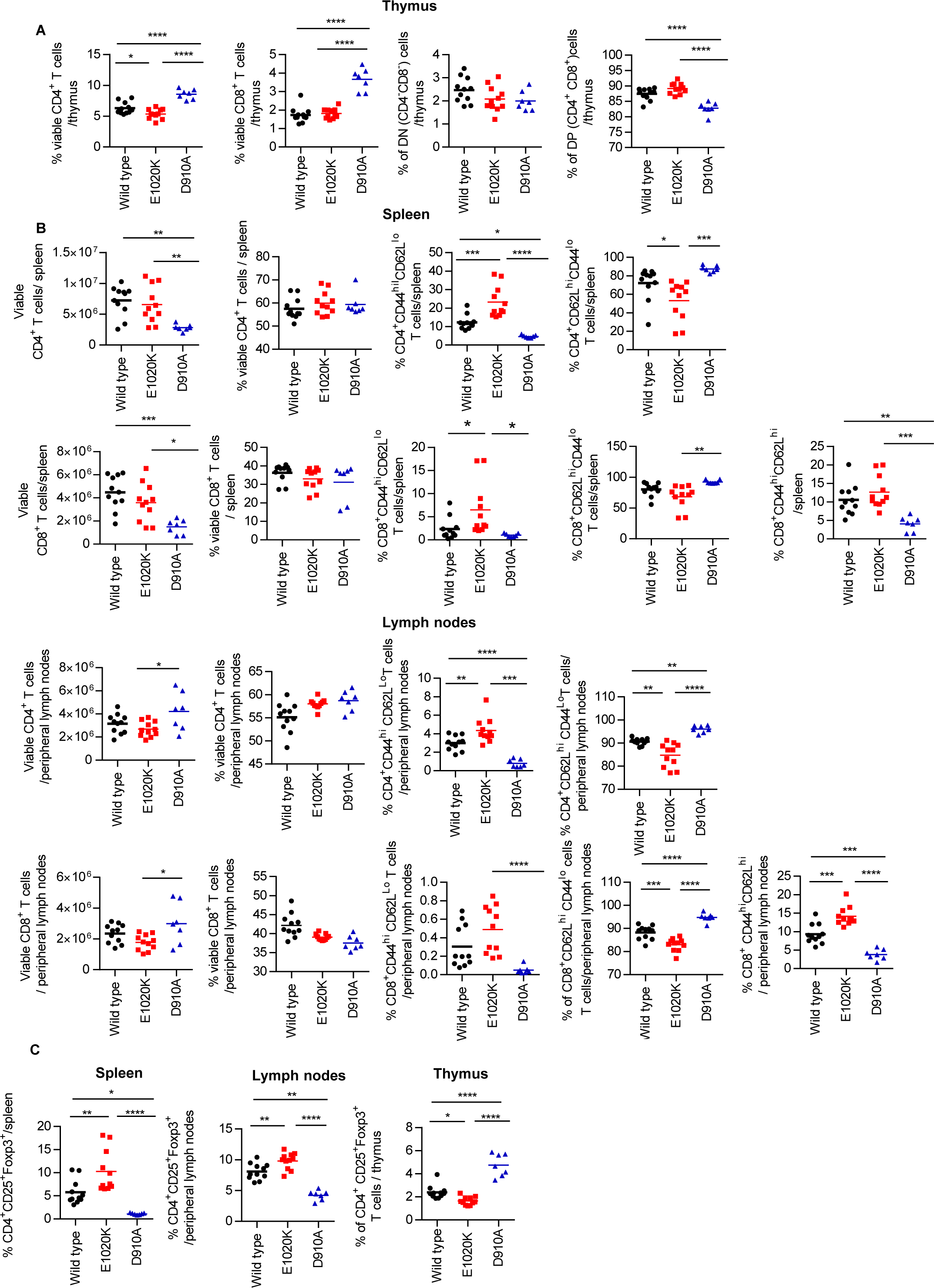
Phenotype of T cells in p110δ^E1020K-T^ mice. T cell populations in the thymus, spleen and lymph nodes in wild-type, p110δ^E1020K-T^ and p110δ^D910A^ were analyzed by flow cytometry. **A:** Thymic T cell proportions in p110δ^E1020K-T^ mice were normal **B:** In the spleen, the proportions of activated CD4^+^ and CD8^+^ T cells (CD44^hi^CD62^lo^) were increased, and there was a trend towards increased proportions of memory CD8^+^ T cells (CD44^hi^CD62L^hi^) in p110δ^E1020K-T^ mice. In the lymph nodes of p110δ^E1020K-T^ mice, the proportions of activated CD4^+^ T cells (CD44^hi^CD62^lo^) were increased with a trend towards increased proportions of activated CD8^+^ T cells, as well as increased proportions of memory CD8^+^ T cells (CD44^hi^CD62L^hi^). **C:** The proportion of CD4^+^CD25^+^Foxp3^+^ regulatory T cells were increased in the lymph nodes and spleen of p110δ^E1020K-T^ mice. (Combined data from 3 independent experiments, n=7-12).

**Supplementary figure 6:**
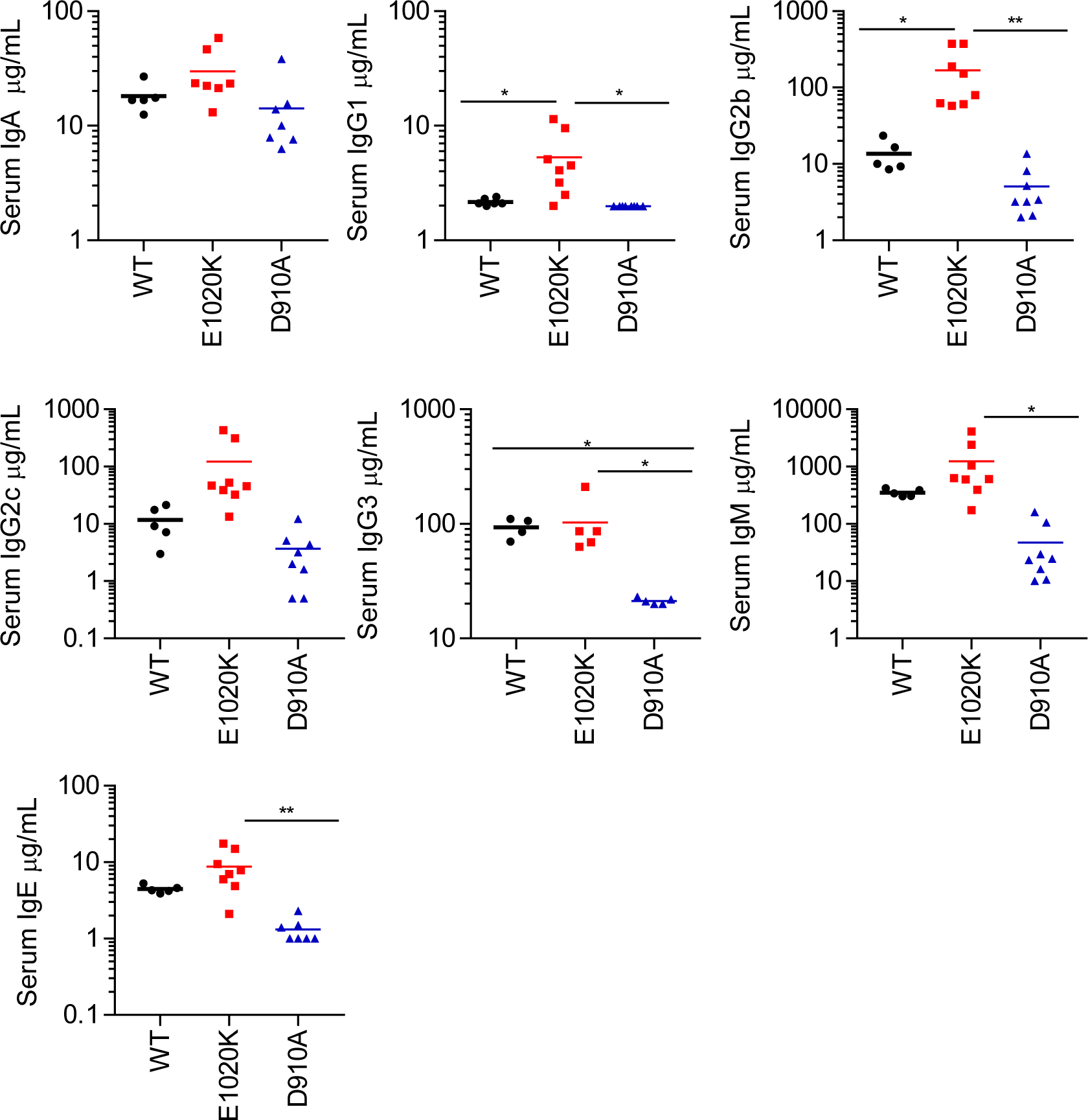
Serum immunoglobulins in p110δ^E1020K-GL^ mice. Analysis of serum immunoglobulins from naïve mice (age 8-12 weeks) showed significantly increased levels of IgG1 and IgG2b, and a trend towards increased levels IgG2c, IgM, IgA, and IgE in p110δ^E1020K-GL^ mice, while IgG3 levels were similar compared to wild-type mice. p110δ^D910A^ mice were antibody deficient for all isotypes analyzed. (n= 6-7).

**Supplementary Figure 7:**
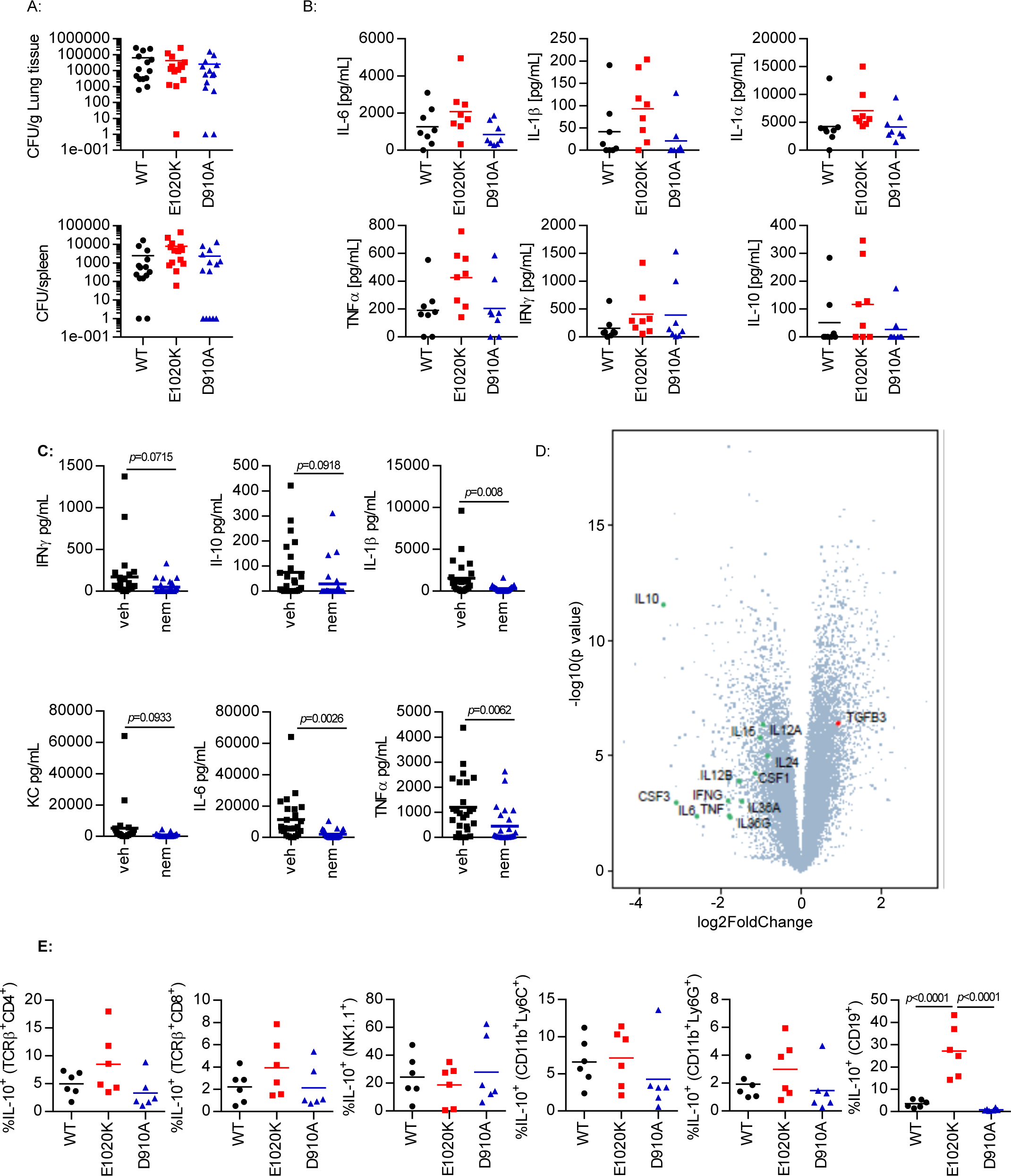
PI3Kδ signaling regulates B cell specific IL-10 production in the lung following *S. pneumoniae* infection. **A:** 24h post *S. pneumoniae* infection, lung and spleen CFU counts were similar in wild-type, p110δ^E1020K-GL^ and p110δ^D910A^ mice. **B:** 24h post infection, cytokine levels in the lung homogenate showed a trend towards increased TNFα, IL-6 and IL-1 in p110δ^E1020K^ mice. **C:** 24h prophylactic treatment with nemiralisib (nem) led to a significant reduction in TNFα, IL-6 and IL-1β, and a trend towards reduced IFNγ and IL-10, in the lungs of wild-type mice compared to vehicle control (veh) treated animals at 24h post infection. **D:** Volcano plot (statistical significance against fold change) of the gene expression changes in response to nemiralisib treatment showed reduced levels of pro-inflammatory cytokines as well as IL-10 at 24h post infection compared to vehicle control treatment. All genes analysed are shown (grey dots) with the cytokines of interest labelled and coloured; green for those with a negative fold change, red for positive fold change. **E:** Analysis of immune cell subsets in *Il10*^ITIB^ reporter mice at 24h post infection showed that the proportion of IL-10 producing B cells is significantly increased in p110δ^E1020K-GL^ mice and reduced in p110δ^D190A^ mice compared to wild-type mice, with similar trends in T cells and myeloid cells not reaching significance. (A: results from 2 independent studies combined, n=14; B: representative data from 3 independent studies n=8; C: Combined data from 4 independent experiments n=25; D: data from 1 study, n=6; E: representative data from 2 independent studies n=6).

**Supplementary Figure 8:**
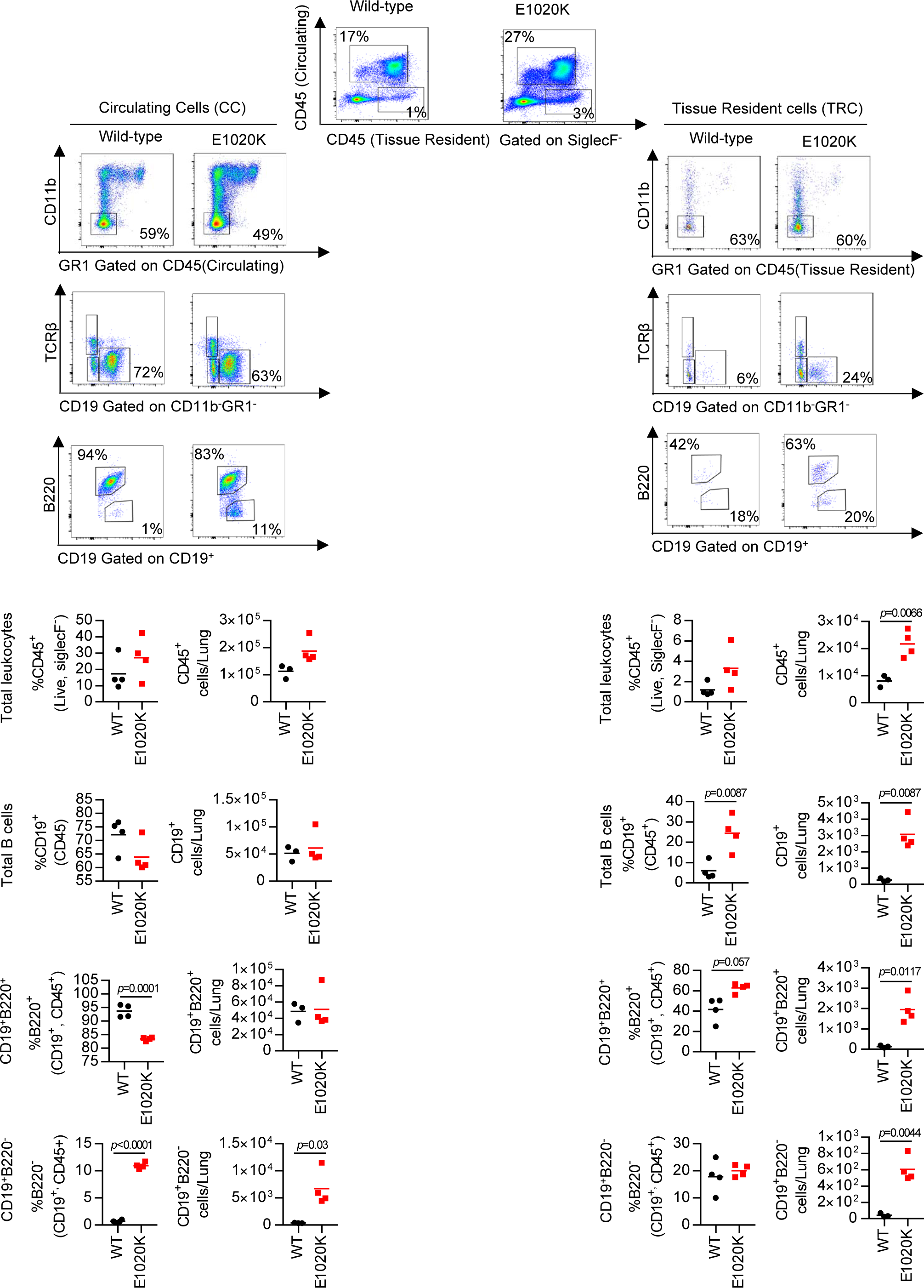
Tissue resident B cell numbers are increased in the lungs of naïve p110δ^E1020K-GL^ mice. Biotin-conjugated anti-CD45 was injected intravenously to distinguish between circulating (CC) and tissue resident (TRC) cells in the lungs of naïve mice. p110δ^E1020K-GL^ mice showed an increase in the proportion and number of tissue resident leukocytes (CD45^+^TRC), but not circulating leukocytes. There was also an increase in the proportion and number of total CD19^+^ B cells among the tissue resident cell population. Among circulating leukocytes (CD45^+^IV), total CD19^+^ B cell numbers were similar in p110δ^WT^ and p110δ^E1020K^ mice, and p110δ^E1020K^ mice had increased numbers and proportions of CD19^+^B220^−^ B cells. (Representative data from 2 independent experiments n=4).

